# DksA, ppGpp and RegAB regulate nitrate respiration in *Paracoccus denitrificans*

**DOI:** 10.1101/2023.01.17.523660

**Authors:** Ashvini Ray, Stephen Spiro

**Affiliations:** Department of Biological Sciences, University of Texas at Dallas, Richardson, Texas 75080

**Keywords:** DksA, guanosine tetraphosphate, nitrate respiration, *Paracoccus denitrificans*, redox homeostasis, RegAB

## Abstract

The periplasmic (NAP) and membrane-associated (Nar) nitrate reductases of *Paracoccus denitrificans* are responsible for nitrate reduction under aerobic and anaerobic conditions, respectively. Expression of NAP is elevated in cells grown on a relatively reduced carbon and energy source (such as butyrate); it is believed that NAP contributes to redox homeostasis by coupling nitrate reduction to the disposal of excess reducing equivalents. Here we show that deletion of either *dksA1* (encoding one of two *dksA* homologs in the *P. denitrificans* genome) or *relA*/*spoT* (encoding a bifunctional ppGpp synthetase and hydrolase) eliminates the butyrate-dependent increase in *nap* promoter and NAP enzyme activity. We conclude that ppGpp likely signals growth on a reduced substrate and, together with DksA1, mediates increased expression of the genes encoding NAP. Support for this model comes from the observation that *nap* promoter activity is increased in cultures exposed to a protein synthesis inhibitor that is known to trigger ppGpp synthesis in other organisms. We also show that, under anaerobic growth conditions, the redox-sensing RegAB two-component pair acts as a negative regulator of NAP expression and as a positive regulator of expression of the membrane-associated nitrate reductase Nar. The *dksA1* and *relA*/*spoT* genes are conditionally synthetically lethal; the double mutant has a null phenotype for growth on butyrate and other reduced substrates while growing normally on succinate and citrate. We also show that the second *dksA* homolog (*dksA2*) and *relA*/*spoT* have roles in regulation of expression of the flavohemoglobin Hmp and in biofilm formation.

**IMPORTANCE:** *Paracoccus denitrificans* is a metabolically versatile Gram-negative bacterium that is used as a model for studies of respiratory metabolism. The organism can utilize nitrate as an electron acceptor for anaerobic respiration, reducing it to dinitrogen via nitrite, nitric oxide, and nitrous oxide. This pathway (known as denitrification) is important as a route for loss of fixed nitrogen from soil and as a source of the greenhouse gas nitrous oxide. Thus, it is important to understand those environmental and genetic factors that govern flux through the denitrification pathway. Here we identify four proteins and a small molecule (ppGpp) which function as previously unknown regulators of expression of enzymes that reduce nitrate and oxidize nitric oxide.

## INTRODUCTION

*Paracoccus denitrificans* is a Gram-negative α-proteobacterium and a facultative anaerobe that can respire using nitrogen oxyanions and oxides as terminal electron acceptors in an anaerobic process known as denitrification (1). In the first step of denitrification, nitrate is reduced to nitrite by the membrane-associated nitrate reductase, Nar (2). *P. denitrificans* also expresses a periplasmic nitrate reductase, NAP, which makes a smaller contribution to the generation of a proton motive force than does Nar (2). Maximal expression of NAP occurs during aerobic growth on relatively reduced substrates, and it is thought that NAP facilitates redox balancing by allowing for the disposal of excess reducing equivalents (3–5).

In the denitrification pathway, the toxic intermediate nitric oxide (NO) is generated from nitrite by the respiratory nitrite reductase and is subsequently reduced to nitrous oxide (N_2_O) by the respiratory NO reductase. *P. denitrificans* also has an *hmp* gene encoding a flavohemoglobin (Hmp) that potentially contributes to the metabolism of NO (6, 7). NO acts in conjunction with transcriptional regulators as an important signal molecule to regulate gene expression during the transition from aerobic respiration to denitrification (8, 9). Pathogenic strains of *Salmonella* and *E. coli* have several transcriptional regulators which control the expression of genes encoding enzymes that scavenge NO. These regulators and their target genes may help to protect the bacteria against the NO that is made by host phagocytes (10–12). One such transcriptional regulator is DksA, which acts as a thiol based NO sensor in *Salmonella* and regulates detoxification of NO through Hmp in *E. coli* (13–15).

DksA, first described in *E. coli* as DnaK suppressor A (16) is a founder member of TraR/DksA family of transcriptional regulators (17, 18). DksA_*Ec*_ (*E. coli* DksA) is 151 amino acids long and consists of three major domains (Fig. S1): a coiled-coil (CC) domain with a conserved DXXDXA motif at its tip, a globular domain with a conserved four-cysteine Zn-finger motif, and a C-terminal α-helical domain (19). The conserved acidic residues located at the tip of the CC domain in DksA_*Ec*_ are critical for binding to RNA polymerase (RNAP) and for DksA_*Ec*_ to function as a transcriptional regulator (20, 21). The globular domain has been shown to interact with the rim of the secondary channel of RNAP (22), and the conserved redox-sensitive cysteines act as a sensor for NO in *Salmonella* (13). Some bacterial genomes encode multiple homologs of DksA, which differ in the presence or absence of a complete DXXDXA motif or the presence of one, two, or four conserved cysteines in the Zn-finger motif (23). For instance, *Pseudomonas aeruginosa* has two functional homologs, DksA1 and DksA2 (24). DksA1 has the conserved four cysteine Zn-finger motif whereas DksA2 does not (25, 26). In α-proteobacteria more closely related to *P. denitrificans*, such as *Sinorhizobium meliloti* and *Rhodobacter sphaeroides*, the assigned *dksA* gene encodes a protein with a complete DXXDXA motif and only one cysteine in the Zn-finger motif (27, 28). Deletion or overexpression of *dksA* in various species has pleiotropic effects on multiple physiological processes. These include amino acid biosynthesis (29), motility (30), biofilm formation (30), plant symbiosis (27), DNA repair (31), quorum sensing and virulence (32) and oxidative and nitrosative damage (13, 15).

DksA, along with the alarmone guanosine-5’, 3’-tetraphosphate (ppGpp) is known to alter global gene expression to adjust cellular metabolism in response to nutritional stress, a mechanism known as the stringent response (when triggered by amino acid starvation) in *E. coli* and other related γ-proteobacteria (33–35). ppGpp is found throughout the bacterial kingdom as a second messenger and is associated with various stress response mechanisms and with growth rate regulation (33). The metabolism of ppGpp is controlled by two enzymes encoded by *relA* and *spoT* (36, 37). RelA is a ribosome-associated ppGpp synthetase, whereas SpoT is a bifunctional enzyme responsible for both synthesis and hydrolysis of ppGpp. The presence of both *relA* and *spoT* is characteristic of γ-proteobacteria, while only one RelA*/*SpoT bifunctional homolog (RSH) possessing both conserved ppGpp synthetase and hydrolase domains is typically present in α-proteobacteria (2, 38). While DksA and ppGpp coregulate many genes, some phenotypic and transcriptomic studies report independent effects of both DksA and ppGpp (39, 40). The independent effects of DksA and ppGpp might be attributed to their direct binding to RNAP; ppGpp is also a direct effector of a number of metabolic enzymes (41–43)

In *P. denitrificans* Pd1222, three gene products have been annotated as members of the TraR/DksA family: Pden_0547 (which we designate DksA1_Pd_), Pden_0916 (DksA2_Pd_), and Pden_1862, which are 45%, 26% and 30% identical to *E. coli* DksA, respectively. The *P. denitrificans* proteins are more similar to their homologs from related α-proteobacteria. DksA1_Pd_, DksA2_Pd_, and Pden_1862 are 90%, 64%, and 33% identical to DksA_Rsp_ (*R. sphaeroides* DksA), respectively, while they are 64%, 47%, and 36% identical to DksA_Sm_ (*S. meliloti* DksA), respectively. *P. denitrificans* also has a single *relA*/*spoT* gene: Pden_1400 (henceforward called *rel*) which encodes for a RelA/SpoT bifunctional homolog (RSH). The predicted RSH is 705 amino acids long and possesses a phosphohydrolase HD domain with a conserved HDXXED motif in the N-terminal region, a RelA-SpoT region, and conserved TGS and ACT domains in the C-terminal domain (Fig. S2). On the basis of sequence similarity, Pden_1400 is predicted to have both ppGpp synthetase and hydrolase activities. None of these *P. denitrificans* genes and proteins has been functionally characterized. A proteomic analysis addressing aerobic to anaerobic growth adaptation in *P. denitrificans* demonstrated a significant increase in expression of DksA1_Pd_ (A1AZG5) under anaerobic conditions. Another study using microarray analysis showed a 3.4-fold higher expression of *dksA2* during anaerobic growth (44, 45). Our attention was drawn to the DksA homologs of *P. denitrificans* because they are differentially expressed under denitrifying conditions, and by the report that DksA regulates *hmp* expression in *E. coli* (15). Our goal for the work described here was therefore to use a reverse-genetics approach in an effort to identify physiological roles for the two DksA proteins and for RSH.

In this work, we report distinct phenotypes associated with deletion of two of the *dksA*-like genes and the *relA/spoT* homolog in *P. denitrificans* Pd1222. We show that deletion of *dksA1* causes an aerobic growth defect on reduced carbon substrates such as butyrate and that *dksA1* and *rel* null strains fail to up-regulate expression and activity of the periplasmic nitrate reductase NAP under these conditions. Furthermore, we demonstrate that a *dksA1 rel* double mutant is unable to grow on reduced carbon substrates. In addition, we show that *dksA2* is involved in biofilm production, possibly indirectly via negative regulation of *hmp* transcription.

## RESULTS

### *P. denitrificans dksA1* and *dksA2* complement Δ*dksA*_*Ec*_ strains

Sequence alignment of the three DksA/TraR proteins revealed that two (Pden_0547 and Pden_0916) have a complete DXXDXA motif which is required for interaction with RNA polymerase (20, 21) while the third (Pden_1862) does not. For this reason, our attention has focused on Pden_0547 and Pden_0916, which we have designated *dksA1* and *dksA2*, respectively. Unlike the canonical DksA of *E. coli*, DksA1_Pd_ has only one cysteine conserved in the zinc finger motif whereas DksA2_Pd_ has all four cysteines (Fig. S1).

*E. coli* MG1655 deleted for *dksA* was complemented with either *dksA1* or *dksA2* cloned in pIND4 and expressed from their native promoter. *E. coli* Δ*dksA* is auxotrophic for multiple amino acids and fails to grow on M9-glucose medium without amino acid supplementation (15). Expression of either *dksA1* or *dksA2* rescued growth of Δ*dksA*_*EC*_ on M9-glucose plates suggesting that either gene may complement the *E. coli dksA* mutation (Fig. 1).

**Figure 1.**
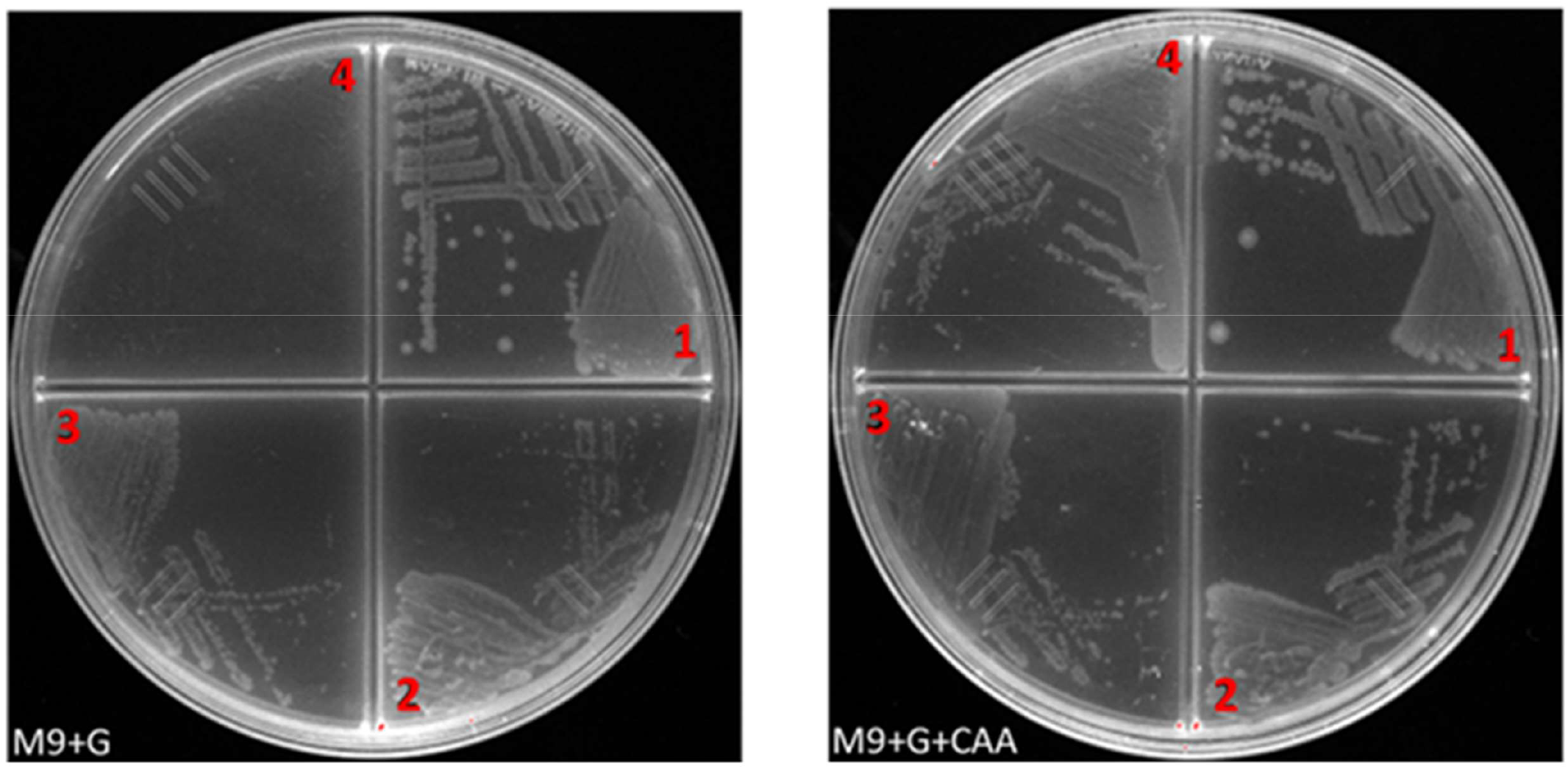
*P. denitrificans dksA1*_*Pd*_ and *dksA2*_*Pd*_ complement an *E. coli* Δ*dksA* mutant (Δ*dksA*_*EC*_). Strains were streaked on M9 minimal (M9) agar plates containing 10 mM glucose (G) with or without 0.2% (w/v) casamino acids (CAA). Plates were incubated overnight at 37 °C. 1 - *E. coli* MG1655; 2 - Δ*dksA*_*EC*_ complemented with *dksA1*_*Pd*_; 3 - Δ*dksA*_*EC*_ complemented with *dksA2*_*Pd*_; 4 - Δ*dksA*_*EC*_ carrying empty pIND4.

### Deletion of *dksA1*_*Pd*_ impairs nitrate stimulation of aerobic growth on butyrate

To characterize the *dksA*-like genes of *P. denitrificans*, we constructed and tested the growth of deletion strains lacking *dksA1* or *dksA2*. Each mutant grew as well as the parent strain on rich (LB) medium under both aerobic and anaerobic conditions (Fig. S3). We also did not observe amino acid auxotrophy when the mutants were grown in succinate or glucose minimal medium (data not shown). In *Salmonella enterica* sv. *typhimurium*, DksA is involved in redox homeostasis since it regulates the expression of enzymes from central metabolic pathways that generate reduced pyridine nucleotides (46). *P. denitrificans* may experience redox imbalance during growth on reduced carbon substrates due to the generation of excess reducing equivalents (3). Therefore, we assayed aerobic growth in minimal medium supplemented with succinate (MM-S) or butyrate (MM-B) as sole carbon and energy sources that are relatively oxidized and reduced, respectively. Growth of the wild-type strain in MM-B was significantly enhanced by the addition of nitrate to the medium. Reduction of nitrate by the periplasmic nitrate reductase NAP provides a mechanism to dispose of excess reducing equivalents during growth on reduced carbon substrates (4). The Δ*dksA1* mutant grew normally on MM-S but showed slightly impaired growth on MM-B relative to the wild type and growth on MM-B was not improved by addition of nitrate to the medium (Fig. 2A and B). Expression of *dksA1* from a recombinant plasmid in the *dksA1* mutant restored a wild type phenotype for growth on MM-B both in the presence and absence of nitrate (Fig. S4). These phenotypes suggest that DksA1_Pd_ may regulate aerobic nitrate reduction by NAP during growth on reduced carbon substrates.

**Figure 2.**
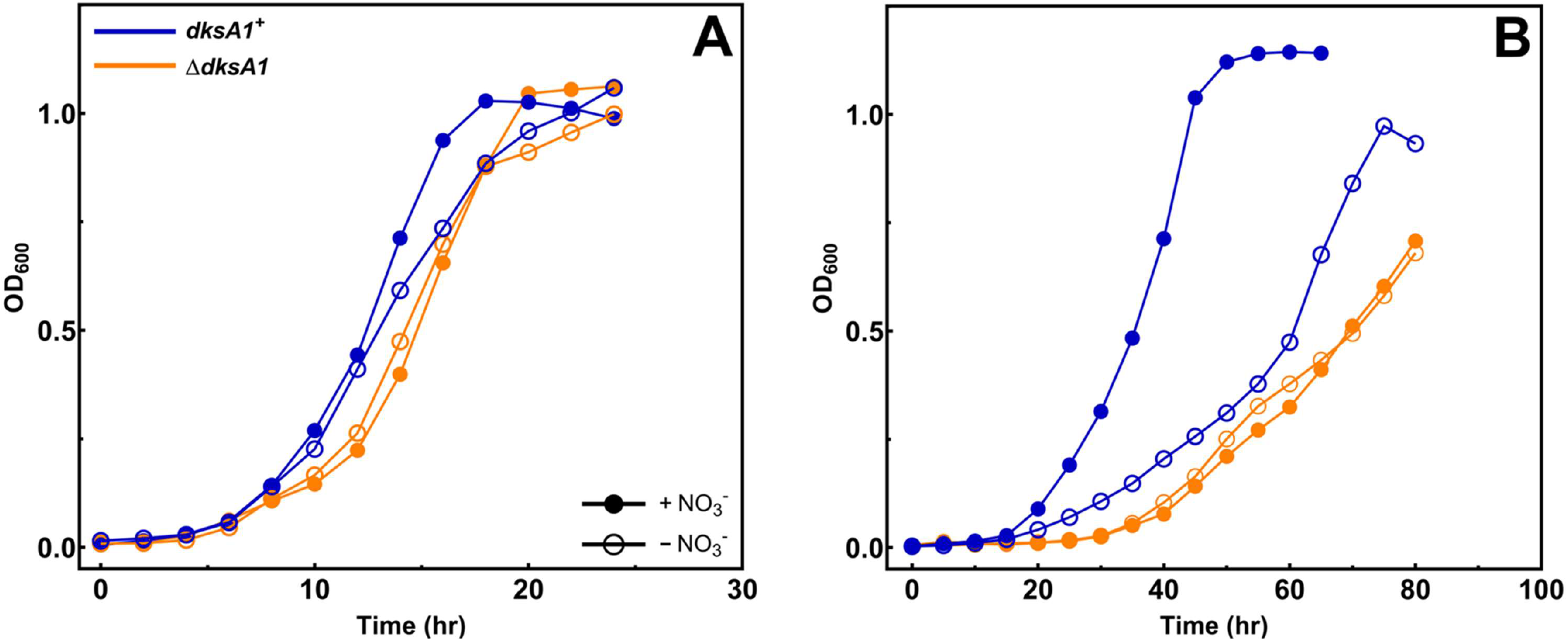
Deletion of *dksA1* impairs nitrate utilization during aerobic growth. Aerobic growth of indicated strains in, **(A)** minimal medium containing succinate or **(B)** minimal medium containing butyrate, with (+ NO_3_^−^) or without (− NO_3_^−^) 20 mM nitrate. Optical density was measured (at 600 nm) at the indicated time points (the means from three independent experiments).

To determine if the Δ*dksA1* associated phenotype is ppGpp-dependent, we measured growth of the Δ*rel* strain under similar conditions. The Δ*rel* mutant showed significantly reduced growth in MM-B compared to wild type, however, unlike for Δ*dksA1*, growth was improved by addition of nitrate to the medium (Fig. 3 and S5), though the magnitude of nitrate stimulation was less than was observed in the wild-type strain. Complementation of the Δ*rel* mutant with the *rel* gene cloned in a plasmid restored the wild type growth phenotypes on MM-B (Fig. S6).

**Figure 3.**
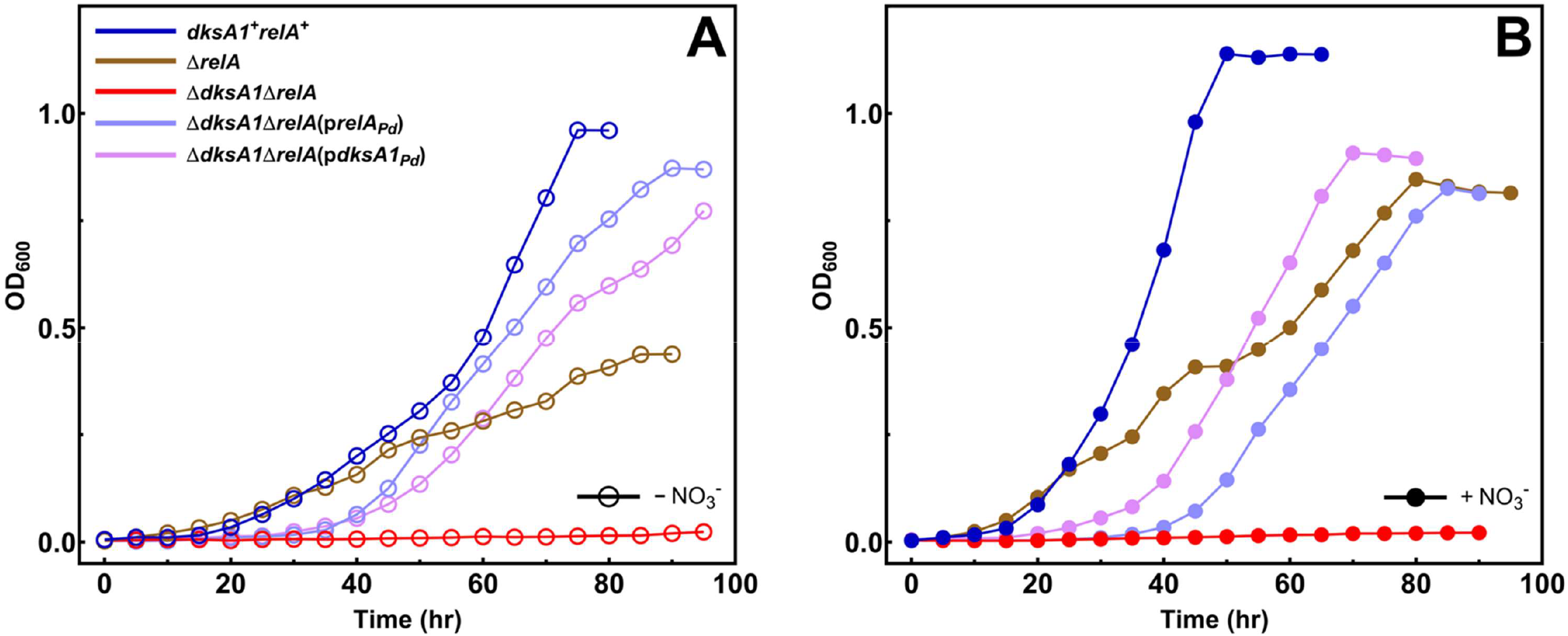
Loss of both *dksA1* and *relA* leads to failure to grow on reduced carbon substrates. Aerobic growth of indicated strains in liquid minimal medium with butyrate **(A)** without (− NO_3_^−^) or **(B)** with 20 mM nitrate (+ NO_3_^−^). Optical density was measured (at 600 nm) at the indicated time points (the means from three independent experiments).

### Δ*dksA1* and Δ*rel* mutants have reduced *napE* promoter activity and NAP enzyme activity

It has been shown previously that the periplasmic nitrate reductase, NAP, has increased expression and activity in cells grown aerobically on reduced energy sources (3, 5). Our results suggest that the absence of *dksA1* impairs nitrate reduction in *P. denitrificans* during aerobic growth. We hypothesized that DksA1_Pd_ regulates expression of NAP during aerobic growth on reduced energy sources. To test this hypothesis, we constructed a *napE-lacZ* promoter transcriptional fusion and measured *napE* promoter activity using β-galactosidase as the reporter. Consistent with previous reports (5), we observed a ∼3-fold increase in *napE* promoter activity in cells grown aerobically in MM-B compared to those grown in MM-S (Fig. 4A). The increase in *napE* promoter activity caused by growth on butyrate was eliminated in the Δ*dksA1* mutant. The increase in *napE* promoter activity in MM-B versus MM-S also disappeared when the cultures were grown anaerobically (Fig. 4A). To confirm that these observations were due to loss of *dksA1*, we measured *napE* promotor activity in the complemented Δ*dksA1* strains. Complementation with the *dksA1* gene restored the butyrate stimulation of *napE* promotor activity (Fig. S7)

**Figure 4.**
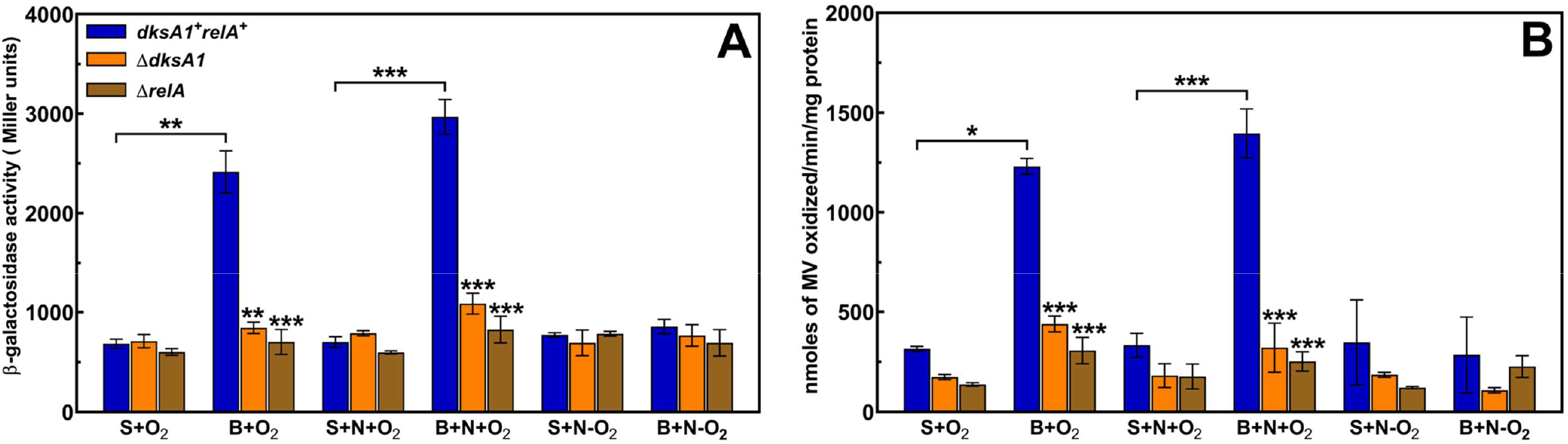
The Δ*dksA1* mutant has reduced *nap* promoter and NAP enzyme activity. **(A)** β-galactosidase activity in *P. denitrificans* wild-type, Δ*dksA1* and Δ*relA* carrying plasmid p*nap*-*lacZ*. **(B)** NAP enzyme activity in the periplasmic fraction of *P. denitrificans* wild-type, Δ*dksA1* and Δ*relA* cells. Cultures were grown in minimal medium containing 10 mM succinate (S) or butyrate (B) as sole carbon and energy source with or without 20 mM nitrate (N). Aerobic (+O_2_) cultures were grown in shake flasks at 250 rpm. Anaerobic (-O_2_) cultures were grown statically in filled closed cap bottles. The bars represent the mean of at least three separate experiments and the error bars represent the standard deviation of the mean. Asterisks on the bars indicate a significant difference compared to the wild-type within the group according to Student’s t-test (*P ≤ 0.05, **P ≤ 0.01, ***P ≤ 0.001).

In parallel with the *napE* promoter activity assays we also measured the activity of NAP in periplasmic fractions using reduced methylviologen as the electron donor to the enzyme. As previously reported (3), and consistent with our promoter activity data, NAP activity in the wild-type was significantly (∼4-fold) higher during aerobic growth in MM-B compared to MM-S (Fig. 4B). The stimulation of activity by growth on butyrate disappeared in the Δ*dksA1* mutants and in cells grown anaerobically in either MM-S or MM-B (Fig. 4B). Our results suggest that DksA1_Pd_ upregulates (either directly or indirectly) the *napE* promoter during aerobic growth on reduced carbon substrates such as butyrate.

The *rel* mutant was found to have the same phenotype as the *dksA1* mutant with respect to *nap* promoter activity and NAP enzyme activity. That is, the butyrate-dependent stimulation was also lost in the *rel* mutant (Fig. 4 and S7). Since DksA and ppGpp frequently act in concert by binding together to a single site on RNA polymerase (34), the likely explanation that DksA-dependent regulation of *nap* requires ppGpp.

### The RegAB system regulates anaerobic *napE* promoter activity and NAP enzyme activity

In *P. pantotrophus*, regulation of the *nap* operon in response to carbon substrate oxidation state involves *cis*-acting regulatory sequences in the *napE* promoter region (48). Specifically, a heptameric imperfect inverted repeat sequence (T<u>G</u>A<u>G</u>A<u>C</u>AtttT<u>GTCGC</u>A) was suggested to be the binding site for a negatively acting redox-responsive regulator (48). The inverted repeat is a partial match to the binding site (GCGGC-N_5_-GTCGC) for the response regulator RegR (RegA) from *Bradyrhizobium japonicum* (49). We found a similar sequence (5′-<u>G</u>A<u>GGC</u>gtttt<u>G</u>A<u>CGC</u>-3′) in the promoter region of the *nap* operon in *P. denitrificans* Pd1222 (matches to the experimentally-determined sequence from *B. japonicum* are underlined). Therefore, we examined *napE* promoter activity in Δ*regA* and Δ*regB* strains to determine if *nap* is regulated by RegAB alongside *dksA1*. In the Δ*regA* mutant, we observed ∼2-fold increases in *napE* promoter activity in cells grown aerobically on succinate (Fig. 5A). In cells grown aerobically on butyrate, *napE* promoter activity was similar in both *regA* and *regB* mutants and the wild-type strain. Interestingly, there were significant increases in promoter activities in *regA* and *regB* mutants grown anaerobically on either succinate or butyrate, compared to the wild type under the same conditions (Fig. 5A). Expression of *regA* and *regB* in the Δ*regA* and Δ*regB* strains respectively reduced *napE* promoter activity to levels below those seen in the wild type strain, confirming that RegAB acts negatively on the *napE* promoter (Fig. S8).

**Figure 5.**
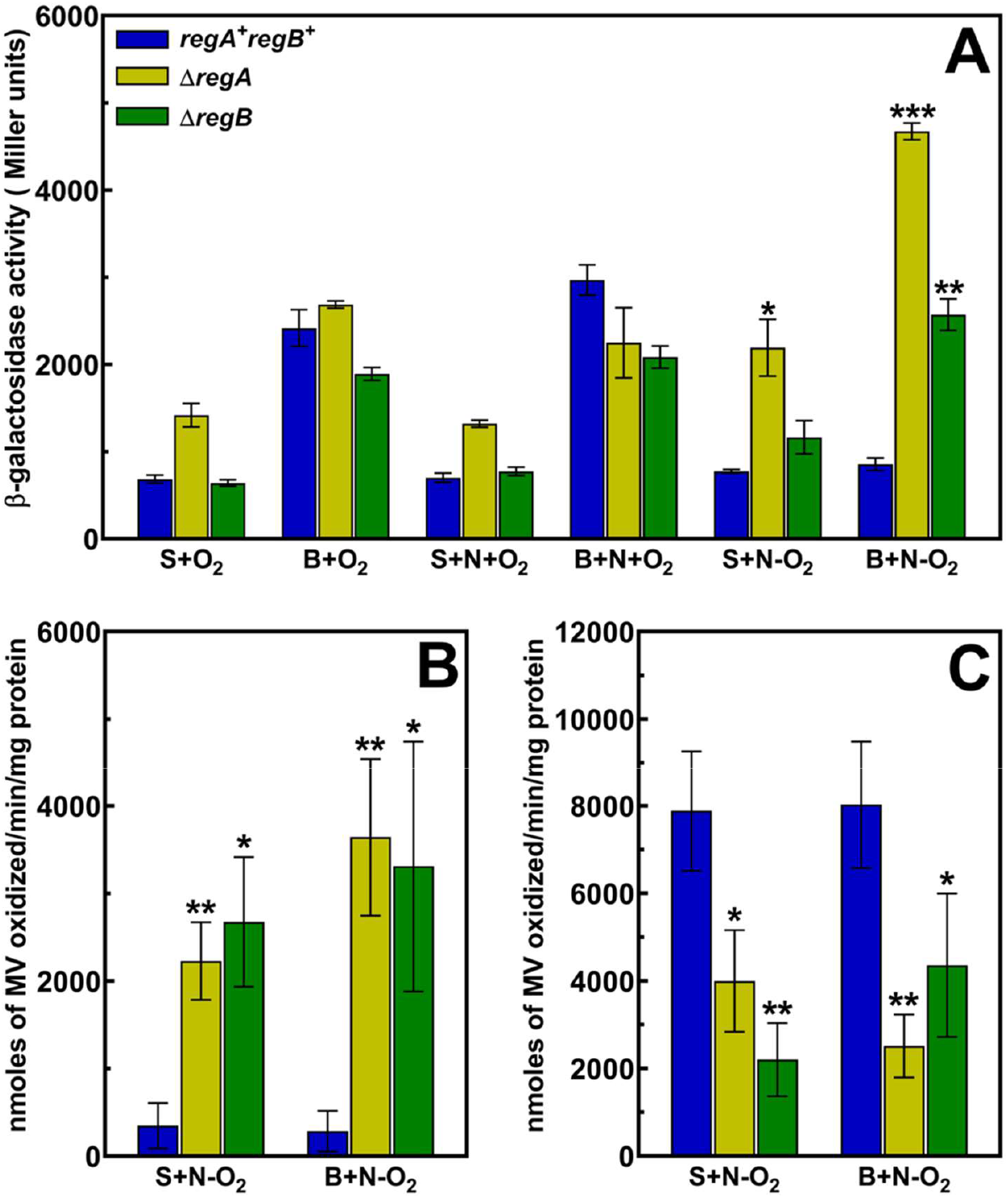
The RegAB system regulates anaerobic *nap* promoter and NAP activity. **(A)** β-galactosidase activity in *P. denitrificans* wild-type, Δ*regA* and Δ*regB* carrying plasmid p*nap*-*lacZ*. **(B)** NAP enzyme activity in the periplasmic fractions of anaerobically grown *P. denitrificans* wild-type, Δ*regA* and Δ*regB* cells. **(C)** NAR enzyme activity in the membrane fractions of *P. denitrificans* wild-type, Δ*regA* and Δ*regB* cells. Cultures were grown in minimal medium containing 10 mM succinate (S) or butyrate (B) as sole carbon source with or without 20 mM nitrate (N). Aerobic (+O_2_) cultures were grown in shake flasks at 250 rpm. Anaerobic (-O_2_) cultures were grown in filled closed cap bottles statically. The bars represent the mean of at least three separate experiments and the error bars represent the standard deviation of the mean. Asterisks on the bars indicate a significant difference compared to the wild-type within the group according to Student’s t-test (*P ≤ 0.05, **P ≤ 0.01, ***P ≤ 0.001).

Our data suggest that RegA is the negatively acting regulator that Ellington et al. suggested (48) binds to the inverted repeat sequence in the *napE* promoter, and that the RegAB system acts to down-regulate this promoter principally under anaerobic growth conditions (but also to a lesser extent during aerobic growth on succinate). The less severe phenotype of the *regB* mutant under some conditions suggests the possibility that RegA is phosphorylated by another histidine kinase (or by a low molecular weight phospho-donor) in the absence of RegB. Assays of NAP activity in periplasmic fractions prepared from anaerobically grown cells confirm that RegAB functions as a negative regulator under these conditions (Fig. 5B).

The membrane-associated nitrate reductase, Nar, is upregulated in *P. denitrificans* in oxygen-limiting conditions (1, 50). Known regulators of the *nar* promoter are NarR (50) and, probably, FnrP (51). Assays of Nar activity in membrane fractions revealed a reduced activity in both Δ*regA* and Δ*regB* mutants suggesting positive regulation of the *nar* operon by RegAB under these conditions (Fig. 5C). Addition of 20 μM sodium azide (a selective inhibitor of Nar) to the reaction mixture inhibited nitrate reduction confirming that the reductase activity in the membrane fraction was due to Nar (data not shown). Reduced Nar activity also coincided with reduced growth yield for the Δ*regA* mutant when grown anaerobically (Fig. S9). Thus, the RegAB system mediates reciprocal regulation of the two nitrate reductases under anaerobic conditions, being a negative regulator of NAP expression and a positive regulator of Nar expression.

### The *dksA1* and *rel* genes are synthetically lethal for growth on reduced substrates

The DksA protein interacts with ppGpp to regulate several functions and deletion of either *dksA* or *rel* gives rise to overlapping pleiotropic phenotypes (39). The similar phenotypes of Δ*dksA1* and *rel* mutants with respect to loss of butyrate stimulation of *nap* promoter activity suggests that DksA1_Pd_ and ppGpp act in concert to regulate expression of the periplasmic nitrate reductase. A Δ*dksA1*Δ*rel* double mutant exhibited a complete growth defect on reduced carbon substrates such as butyrate, caproate and choline (Fig. 3, Fig. S10), while growth on the relatively oxidized substrates succinate and citrate was normal (Fig. S11). Expression of either *dksA1* or *rel* on a broad host range plasmid from their native promoters restored growth of the Δ*dksA1*Δ*rel* mutant, confirming that both mutations are required for the null growth phenotype (Fig. S12). Thus, *dksA1* and *rel* exhibit synthetic lethality with respect to growth on reduced substrates. The likely explanation is that two (or more) genes that are independently regulated by DksA1_Pd_ and ppGpp are together required for growth on reduced substrates.

### Partial activation of the *napE* promoter by a protein synthesis inhibitor

In *E. coli*, ppGpp synthesis by RelA can be stimulated by manipulations that reduce the extent of aminoacyl tRNA charging, including amino acid limitation or inhibition of an amino acyl tRNA synthetase (52). For example, serine hydroxamate (SHX) is a seryl-tRNA synthetase inhibitor that triggers ppGpp synthesis (52–54). SHX (1.28 mM) inhibits *E. coli* growth, and inhibition can be reversed by exogenous serine, presumably because serine overcomes competitive inhibition mediated by SHX (53). Addition of 3 mM SHX to MM-S caused a prolonged growth inhibition of wild type *P. denitrificans* that was reversed by the addition of casamino acids to the medium (Fig. 6). The physiological basis for the eventual recovery of the culture is not known but it may be ppGpp-dependent since the *rel* mutant did not recover from the SHX treatment (Fig. 6). The simplest interpretation of these observations is that SHX treatment triggers ppGpp synthesis by RSH and that ppGpp is required for the recovery from growth arrest. A prediction that follows is that treatment of a culture with a sub-inhibitory concentration of SHX will stimulate a promoter activated by ppGpp. Therefore, we measured activity of the *napE*-*lacZ* fusion in cultures grown aerobically in MM-S and treated with 2 mM SHX while in the logarithmic phase of growth. Ninety minutes after SHX addition, promoter activity in the treated cultures was ∼2-fold higher than in the untreated cultures (1101 ± 96 versus 609 ± 60 units of β-galactosidase activity). In the *rel* mutant, we saw no induction of the *napE* promoter in response to 2 mM SHX. Partial induction of the *napE* promoter by a treatment known to trigger ppGpp synthesis in other organisms is consistent with the idea that the promoter is regulated by ppGpp and DksA1.

**Figure 6.**
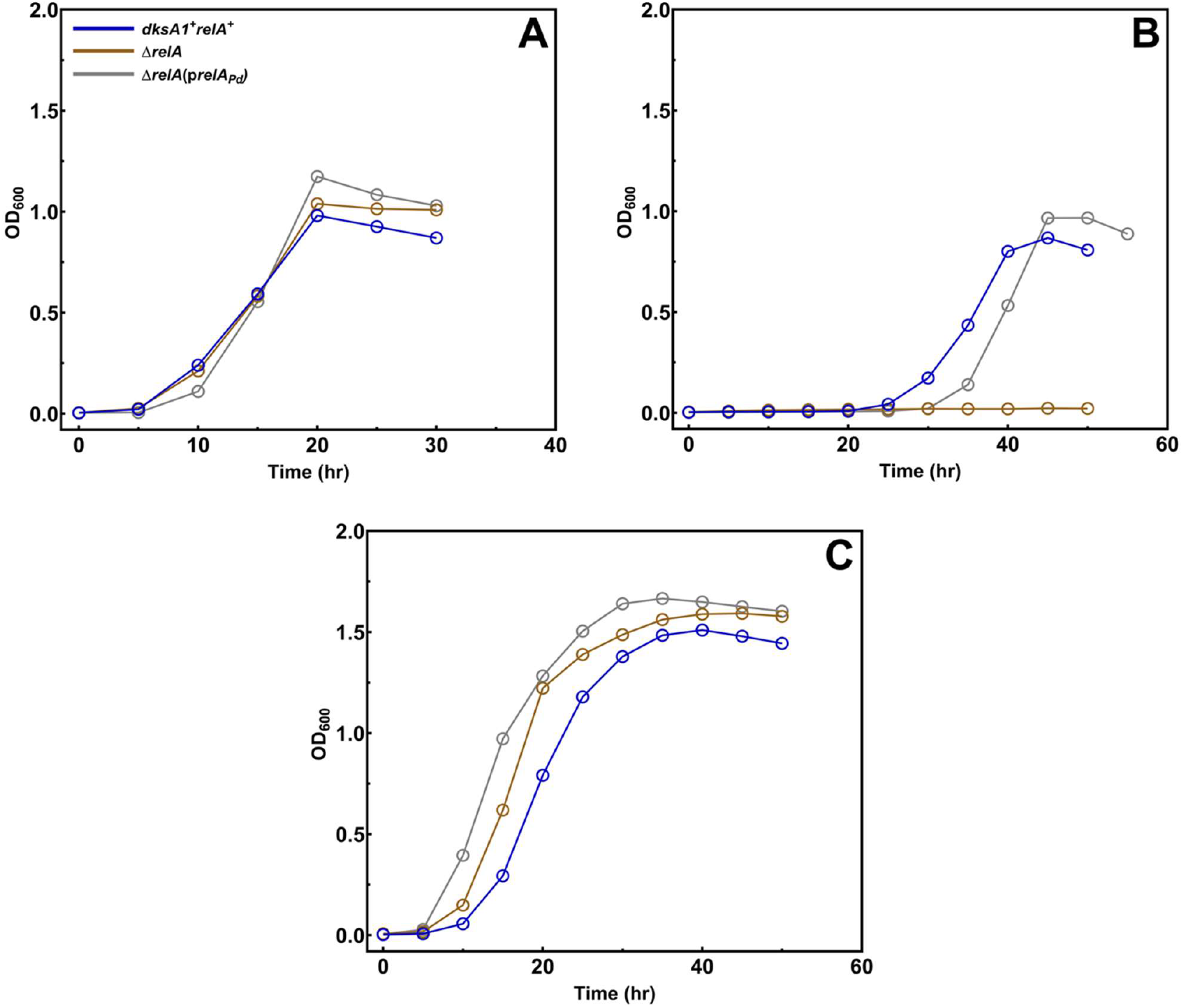
The *relA* gene is required for the response to amino acid starvation in *P. denitrificans*. Aerobic growth of indicated strains in minimal medium containing: **(A)** succinate only, **(B)** succinate and 3 mM serine hydroxamate, and **(C)** succinate, 3 mM serine hydroxamate and 0.2% (w/v) casamino acids. Optical density was measured (at 600 nm) at the indicated time points (the means from three independent experiments).

### DksA2_Pd_ is a positive regulator of the *hmp* promoter

In *E. coli, dksA* regulates the expression and activity of Hmp, a flavohemoprotein. Hmp is a nitric oxide dioxygenase or denitrosylase which oxidizes nitric oxide to nitrate, and is known to be the dominant NO defense mechanism under aerobic conditions (15). *P. denitrificans* has an *hmp* gene, Pden_1689 (6, 7) and we have shown that the Hmp is functional for NO scavenging (unpublished data). To determine if regulation of *hmp* is mediated by DksA in this organism, we assayed *hmp* promoter activity in *P. denitrificans* and the Δ*dksA* mutants. Our results indicate that *hmp* promoter activity is modestly but significantly elevated in the Δ*dksA2* mutant but is unaffected by the Δ*dksA1* deletion (Fig. 7A). The Δ*rel* mutant also shows higher *hmp* promoter activity similar to the Δ*dksA2* strain suggesting that the regulation of *hmp* by DksA2 is ppGpp-dependent (Fig. 7A).

**Figure 7.**
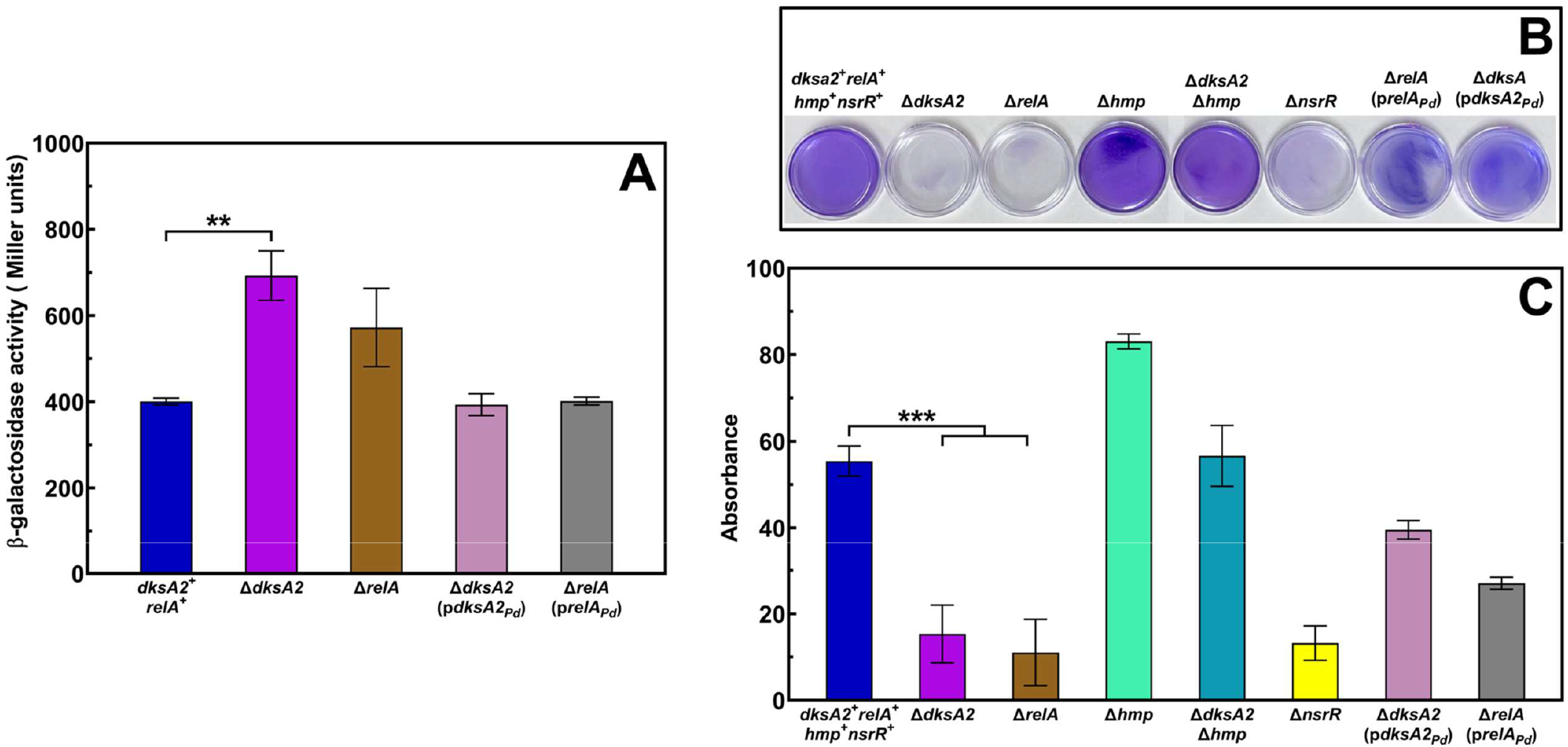
DksA2_Pd_ and RSH regulate *hmp* promoter activity and biofilm formation in *P. denitrificans*. **(A)** β-galactosidase activities in *P. denitrificans* wild-type, Δ*dksA2* and Δ*relA* carrying plasmid p*hmp*-*lacZ*. **(B)** Assays of biofilm formation by the wild-type, deletion, and complemented strains. Dishes were washed, stained with crystal violet, and photographed from above. **(C)** The crystal violet was extracted with 95% ethanol, and the absorbance measured at 595 nm. Means and standard deviations were calculated from results of three separate assays. Cultures were grown in 6-cm-diameter petri dishes in liquid L broth (Miller), supplemented with 10 mM CaCl_2_ and incubated without shaking at 30 °C. Cells were harvested at mid-log phase for the β-galactosidase assay or left undisturbed for 72 hours for the biofilm assay. Asterisks on the bars indicate a significant difference compared to the wild-type according to Student’s t-test (**P ≤ 0.01, ***P ≤ 0.001).

### Both DksA2_Pd_ and RSH regulate biofilm formation in *P. denitrificans*

Our previous work has shown that deletion of *hmp* increases biofilm formation in *P. denitrificans* Pd1222 possibly due to accumulation of NO (6). Given the evidence that DksA2_Pd_ regulates *hmp* expression in *P. denitrificans*, we assayed attached growth to determine if DksA2_Pd_ also affects biofilm formation. We observed that Δ*dksA2* strains had greatly reduced levels of attached growth (Fig. 7B and C). Addition of an *hmp* mutation to the *dksA2* mutant rescued biofilm formation to wild-type levels (Fig. 7B and C) suggesting that the *dksA2* phenotype requires Hmp. Deletion of *nsrR*, which encodes a transcriptional repressor of *hmp* (6), led to significantly reduced biofilm formation similar to Δ*dksA2* (Fig. 7B and C). We also observed reduced biofilm in a Δ*rel* strain (and in a Δ*dksA2* Δ*rel* double mutant, not shown) suggesting that *dksA2* and *rel* could be jointly involved in regulating biofilm formation in *P. denitrificans* via negative regulation of *hmp* (Fig. 7B and C).

## DISCUSSION

Expression of the periplasmic nitrate reductase NAP is upregulated by aerobic growth on reduced carbon and energy sources, the likely explanation being that nitrate reduction by NAP provides a means to dispose of excess reducing equivalents (3–5, 47, 48). We have shown that DksA1_Pd_ and RSH are required for the butyrate-stimulation of *napE* promoter and NAP enzyme activity. RSH is the only protein encoded in the *P. denitrificans* genome with predicted ppGpp synthetase activity and that it is enzymatically functional is supported by the detection of ppGpp in slow-growing cultures of this organism (55). It is likely that DksA1_Pd_ and ppGpp directly activate the *napE* promoter, although indirect effects cannot be excluded. In either case, the implication is that growth on reduced substrates somehow triggers the synthesis of ppGpp by RSH. Data from continuous cultures show that the *napE* promoter is regulated both by growth rate and by carbon substrate, leading to the conclusion that “…*nap* regulation is affected by a physiological signal which integrates aspects of growth rate and carbon substrate utilization” (56). Our work suggests that ppGpp (together with DksA1_Pd_) is the integrating signal, which is consistent with the well-known role of ppGpp in growth rate control (33).

One difference between the phenotypes of the *dksA1* and *rel* mutants is that nitrate does not improve aerobic growth of the *dksA1* mutant on butyrate (as it does in the case of the wild-type strain) but does modestly improve growth of the *rel* mutant. This improvement of growth cannot be attributed to butyrate stimulation of *nap* expression, since that is lost in the *rel* mutant. It is also the case that the *rel* mutant grows less well on butyrate (in the absence of nitrate) than the wild-type strain and the *dksA1* mutant, and that the double mutant does not grow at all on reduced substrates. It is likely that there are DksA1_Pd_ and ppGpp regulated genes that are involved in redox homeostasis that are yet to be identified.

We showed that regulation of *nap* expression also involves the RegAB two-component system, which is redox-sensing and involved in redox homeostasis (57). Our results show that RegAB is not a major player in mediating up-regulation of the *nap* promoter by growth on butyrate, but instead negatively regulates *nap* expression during anaerobic growth. The response regulator RegA very likely binds to an inverted repeat in the *napE* promoter that was initially characterized in the closely related organism *P. pantotrophus* and suggested to be the binding site for a negatively acting regulator (48). Evidently, regulation of the *napE* promoter by carbon substrate and by oxygen availability can be separated, with the former requiring DksA1_Pd_ and ppGpp and the latter RegAB. Interestingly, RegAB also contribute (either directly or indirectly) to the upregulation of the membrane associated nitrate reductase under anaerobic growth conditions. Thus, NAP and Nar are reciprocally regulated by oxygen availability to be the dominant nitrate reductases under aerobic and anaerobic conditions, respectively.

A recent study has shown that RegAB acts both positively and negatively under hypoxic conditions in *Burkholderia pseudomallei*, an α-proteobacterium quite closely related to *P. denitrificans* (58). Specifically, RegAB are required for upregulation of Nar under hypoxic conditions in this organism, though this may involve both direct regulation by RegA and indirect regulation, via FNR (58). Indirect effects of RegAB via NarR and/or FnrP are also possible in *P. denitrificans*; we are able to discern a potential RegA binding site in the *narR*-*narK* intergenic region that controls expression of the *narGHJI* genes encoding Nar (50).

In *E. coli*, DksA and ppGpp contribute to regulation of the *hmp* gene encoding the flavohemoglobin, Hmp, which is responsible for detoxifying nitric oxide (15, 59, 60). In *P. denitrificans*, we find that deletion of *dksA2* or *rel* leads to modestly increased *hmp* promoter activity and dramatically reduced ability to form biofilm. Since the biofilm-defective phenotype requires Hmp, we suggest that reduced expression of *hmp* in the mutants causes a reduced accumulation of nitric oxide, which stimulates biofilm formation in this organism (6). Reduced growth as biofilm in *dksA* mutants has been reported for other organisms, though the regulatory mechanisms are likely different (30, 61).

Our study has identified two functional DksA proteins in *P. denitrificans* that have distinct roles in growth on reduced carbon substrates, aerobic and anaerobic nitrate respiration, expression of flavohemoglobin, and biofilm formation. In other α-proteobacteria which have at least two DksA homologs, physiological roles have so far been ascribed to only one of them (27, 28). Further functional analysis of the *P. denitrificans* DksA-like proteins and of the role of ppGpp will advance our broader understanding of DksA functions in α-proteobacteria.

## MATERIALS AND METHODS

### Bacterial strains and growth conditions

The bacterial strains and plasmids used are listed in Table S1. All *Paracoccus denitrificans* strains were derived from the wild-type Pd1222 (62). *Escherichia coli* S.17-1 was used as the donor and *E. coli* DH5α (pRK2013) as the helper strain for the introduction of plasmids into *P. denitrificans* by conjugation (63, 64). *E. coli* Novablue competent cells (Novagen) were used as the host for all genetic manipulations. The rich medium used for routine growth was LB (Miller) Broth (10 g/l tryptone, 5 g/l yeast extract, 10 g/l NaCl), and the defined minimal medium (MM) for *P. denitrificans* was as previously described (65), solidified with 1.5 % agar (w/v) as needed and supplemented with carbon and energy sources and with 20 mM nitrate as indicated. For growth assays, *P. denitrificans* strains were first streaked on LB agar plates, single colonies were inoculated into LB broth and grown overnight in 15 ml tubes at 250 rpm. These cells were washed with MM, inoculated into succinate-supplemented MM, and grown overnight in 15 ml tubes at 250 rpm. Cells were pelleted, washed, and finally inoculated into MM supplemented with different carbon sources with or without nitrate. For aerobic growth assays, cultures were grown in triplicate wells in a BioTek plate reader with horizontal orbital shaking at 280 rpm and a volume of 0.2 ml (<50% of total volume) to ensure proper aeration while preventing cross contamination. The optical density was measured at 600 nm and adjusted for path length variation. Serine hydroxamate was added directly from a freshly made 100 mM stock to the growth medium to a final concentration of 2-3 mM either at the time of inoculation or to a growing culture, as indicated.

Anaerobic cultures for β-galactosidase assays were grown in filled glass cuvettes sealed with a cap and septum, with medium supplemented with 20 mM nitrate. The cuvettes were gassed with N_2_ and incubated statically. Aerobic cultures were grown in 250 ml flasks shaking at 250 rpm in MM supplemented with the indicated carbon substrate and with or without 20 mM nitrate. For *E. coli* complementation assays, solid M9 medium (66) with 10 mM glucose was used, with or without 0.2 % (w/v) casamino acids.

Plasmids were maintained by addition of antibiotics at the following concentrations: ampicillin 100 μg/ml, kanamycin 25 μg/ml, rifampicin 100 μg/ml, tetracycline 2 μg/ml for *P. denitrificans* and 15 μg/ml for *E. coli*. All *P. denitrificans* strains were grown at 30 °C and *E. coli* at 37 °C.

### Genetic manipulations

All primers used are listed in Table S2. Unmarked deletions were made by allelic replacement using pK18*mobsacB* as described by Sullivan et al. with minor modifications (67, 68). Briefly, in-frame marker-less deletions were constructed by PCR amplification of ∼500-1000 bp fragments from either side of the gene of interest. The two amplicons were ligated together either by PCR using overlapping regions added to the primers or by digestion and ligation using a restriction site incorporated in the primers (69). The ligated fragment was first cloned into pSTBlue-1 (Novagen), then sub-cloned into pK18*mobsacB*, transformed into *E. coli* S.17-1 and mobilized into *P. denitrificans* by conjugation. Triparental mating was performed by mixing *P. denitrificans, E. coli* S.17-1 harboring recombinant pK18*mobsacB* or pMP220 or pIND4 derivatives, and *E. coli* DH5α (pRK2013) cells grown to log phase, in a 2:1:1 ratio and by incubation on LB agar plate for 2-3 days at 30 °C. Single-crossover recombination events were selected on rifampicin and kanamycin plates, and then recombinants with double crossovers were isolated by growth on 6 % (w/v) sucrose plates and kanamycin sensitivity as described by Schafer et al. (67). All mutations were designed to be in-frame deletions to prevent polarity. Promoter-*lacZ* reporter fusions were constructed by amplifying ∼300-400 bp upstream of the start codon of genes of interest and first cloning into pSTBlue-1. The sequence was verified and the fragment cloned into the low copy number vector pMP220 between *Eco*RI and *Pst*I restriction sites (70). Plasmids were introduced into *P. denitrificans* by conjugation from *E. coli* S.17-1. *P. denitrificans* exconjugants were selected by rifampicin and tetracycline resistance. For complementation tests in *P. denitrificans*, the gene of interest with its native promoter was amplified and cloned directly into pIND4 (71) and the sequence verified. The cloning was done using the NEBuilder HiFi DNA assembly cloning kit (NEB). Plasmids were transformed into *E. coli* S.17-1 and mobilized into *P. denitrificans* strains by conjugation as described above.

### β-galactosidase assay

Liquid cultures were grown aerobically and anaerobically in minimal medium with the indicated carbon sources and with or without 20 mM nitrate. Cells were harvested at the mid-log phase of growth and the assay was performed as described by Miller (66). β-galactosidase activity was measured using 0.1 ml of cells made permeable by treatment with sodium dodecyl sulfate and chloroform. In the case of the *hmp-lacZ* fusion, strains were grown to mid-log phase in the same medium and growth conditions as used for the biofilm assays.

### Biofilm assays

Attached growth was assayed in 60 mm × 15 mm polystyrene petri dishes (6). A 70 μl aliquot of overnight culture was inoculated into 7 ml of LB broth supplemented with 10 mM calcium chloride. The Petri dishes were incubated statically at 30 °C for 72 hours. The plates were then washed thrice under gently running tap water and stained with crystal violet (0.1 % in water) for 20 minutes, washed three times with water, and air dried. For quantitation, the crystal violet stain was extracted into 3 ml 95 % ethanol, diluted as necessary, and absorbance measured at 595 nm (6, 72).

### Cell fractionation

*P. denitrificans* strains were grown aerobically and anaerobically in MM with either succinate (10 mM) or butyrate (10 mM) as the sole carbon source; nitrate (20 mM) was added as indicated. Aerobic cultures (50 ml) were grown at 30 °C in 250 ml conical flasks shaken at 250 rpm. Anaerobic cultures were grown in 100 ml bottles filled to the neck with medium and incubated at 30 °C without shaking. Periplasmic fractions for the nitrate reductase assay were prepared as described in Quan et al (73). In short, mid-exponential phase cells were harvested by centrifugation at 3,000 *g* for 20 minutes at 4 °C. Cells were then gently resuspended in TSE buffer (200 mM Tris–HCl, pH 8.0, 500 mM sucrose, 1 mM EDTA without any protease inhibitor) and incubated on ice for 30 minutes, and centrifuged at 16,000 *g* for 30 minutes at 4 °C. The supernatant was transferred to a fresh tube and used as the periplasmic fraction. Membrane fractions were prepared according to Dunn et al. with some modifications (74). Briefly, cultures were harvested in the mid-exponential phase of growth by centrifugation at 8,000 *g* for 5 minutes at 4 °C and washed twice in 10 mM potassium phosphate buffer (pH 7.5) in 50 ml Falcon tubes. Cells were resuspended in 2.5 ml of ice cold 10 mM phosphate buffer, 10 mM MgSO_4_, 0.2 M NaCl. Samples were sonicated on ice using a Fisher Scientific Sonic Dismembrator model 500 outfitted with a 3 mm diameter tip at 20 % amplitude (10 W) for five 20 second bursts at 1 minute intervals. To clarify the lysate, samples were centrifuged at 16,000 *g* for 20 minutes at 4 °C. The clarified total lysate was further centrifuged at 100,000 *g* for 2 hours at 4 °C. The pellet was suspended in 0.2 ml of phosphate buffer and retained as the membrane fraction for the enzyme assay. Fractions were kept on ice and used within hours of preparation.

### Enzyme activity

Nitrate reductase activity was determined spectrophotometrically using reduced methylviologen (MV^+^) as the electron donor and 10 mM sodium nitrate as the electron acceptor (3). The extinction coefficient of reduced methylviologen used was 8.25 mM^−1^ cm^−1^ at 600 nm (75). The protein content of the periplasmic and membrane fractions was measured by the protein estimation method of Lowry et al., using bovine serum albumin as the standard (76).

## ACKNOWLEDGMENTS

We are grateful to Victor Luque-Almagro, Andrew Gates and David Richardson for generously providing the *regA*::*kan* mutant and to Aashna Pathi for her contribution to strain construction.

