## Supplemental Figures 1-12 for "DksA, ppGpp and RegAB regulate nitrate respiration in *Paracoccus denitrificans*"

**Supplementary figures**

### Fig. S1

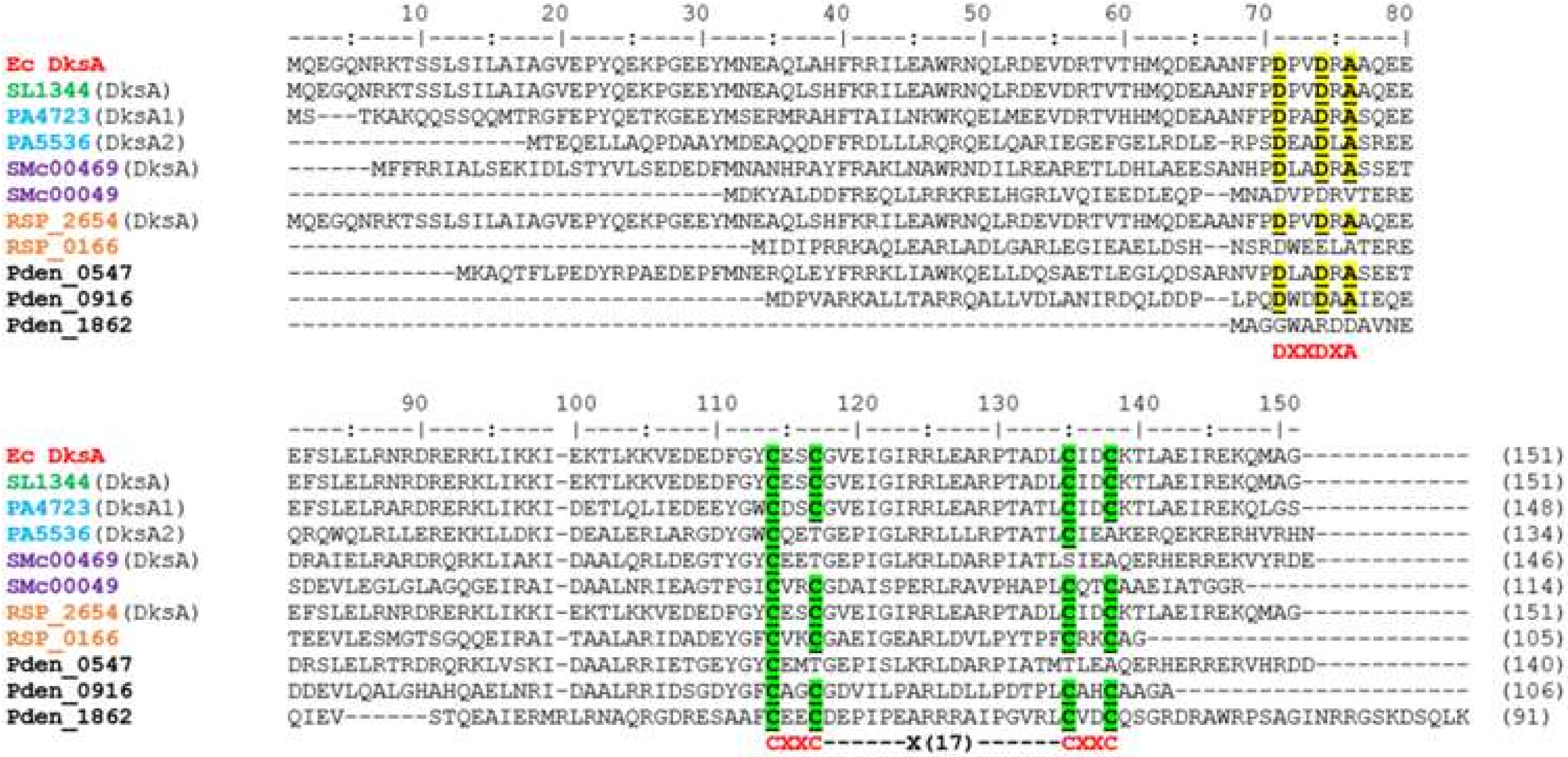


**Figure S1. Alignment of various DksA homologs.** Bacterial genomes encode one or more DksA homologs, differing in the presence or absence of a DXXDXA motif (highlighted in yellow), or in the number of conserved cysteines (1, 2 or 4) within the zinc finger motif (green). Amino acid sequences (from NCBI) were aligned using ClustalW. The numbers on the top of alignment represent residue position of *E. coli* DksA. Ec, *Escherichia coli*; SL, *Salmonella enterica*; PA, *Pseudomonas aeruginosa*; SMc, *Sinorhizobium meliloti*; RSP, *Rhodobacter sphaeroides*; Pden, *Paracoccus denitrificans*.

### Fig. S2

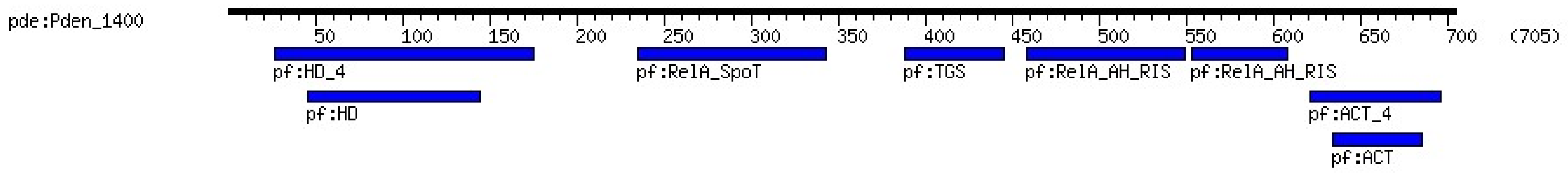


**Figure S2. RelA/SpoT (RSH) homolog of *Paracoccus denitrificans*.** Protein sequence analysis from the KEGG database shows that the protein encoded by *Pden_1400* has domain organization similar to RelA/SpoT bifunctional proteins. Shown in the figure are the N-terminal conserved phosphohydrolase HD domain, a predicted RelA-SpoT region, and the C-terminal TGS and ACT regulatory domains.

<u>**Supplementary figures**</u>

### Fig. S3

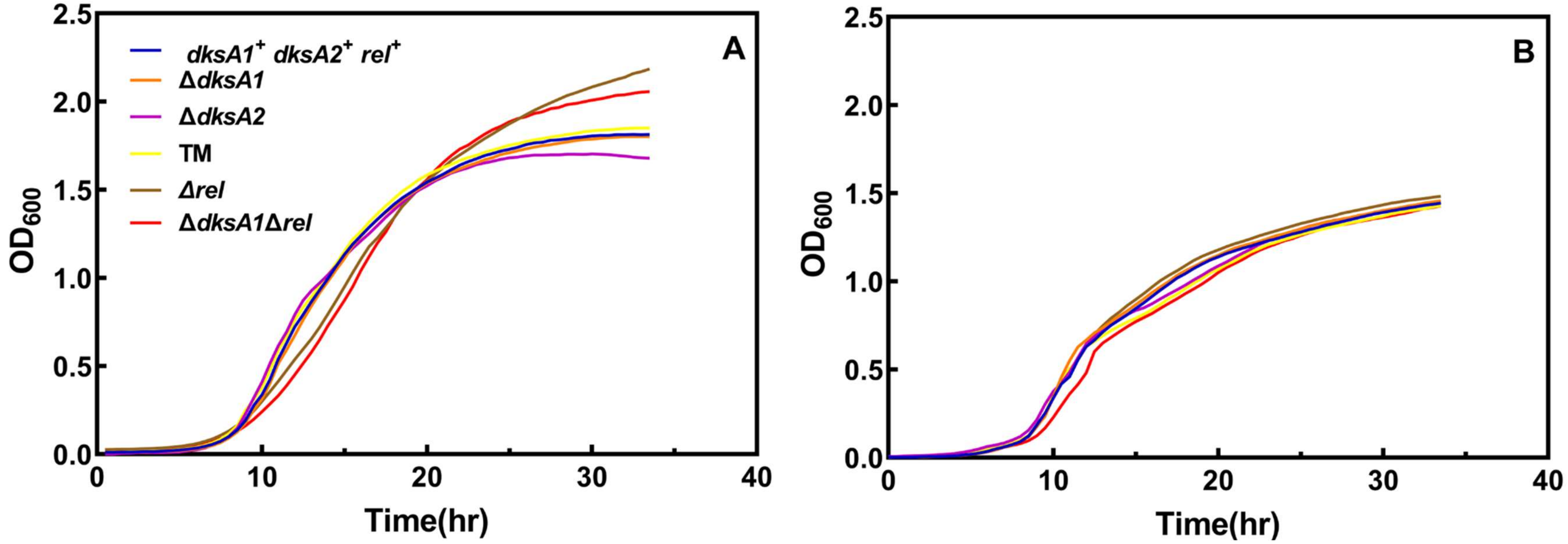


**Figure S3. *dksA* and *relA* deficient strains grow same as WT in LB/rich medium. (A)** aerobic and **(B)** anaerobic growth of wild-type *P. denitrificans*, Δ*dksA1*, Δ*dksA2*, TM (Δ*dksA1*Δ*dksA2*Δ*Pden_1862*), Δ*rel*, and Δ*dksA1*Δ*rel*, in liquid LB. Strains were grown in liquid L broth (Miller), aerobically and anaerobically at 30 ºC. The solid line represents mean of three independent replicates.

### Fig. S4

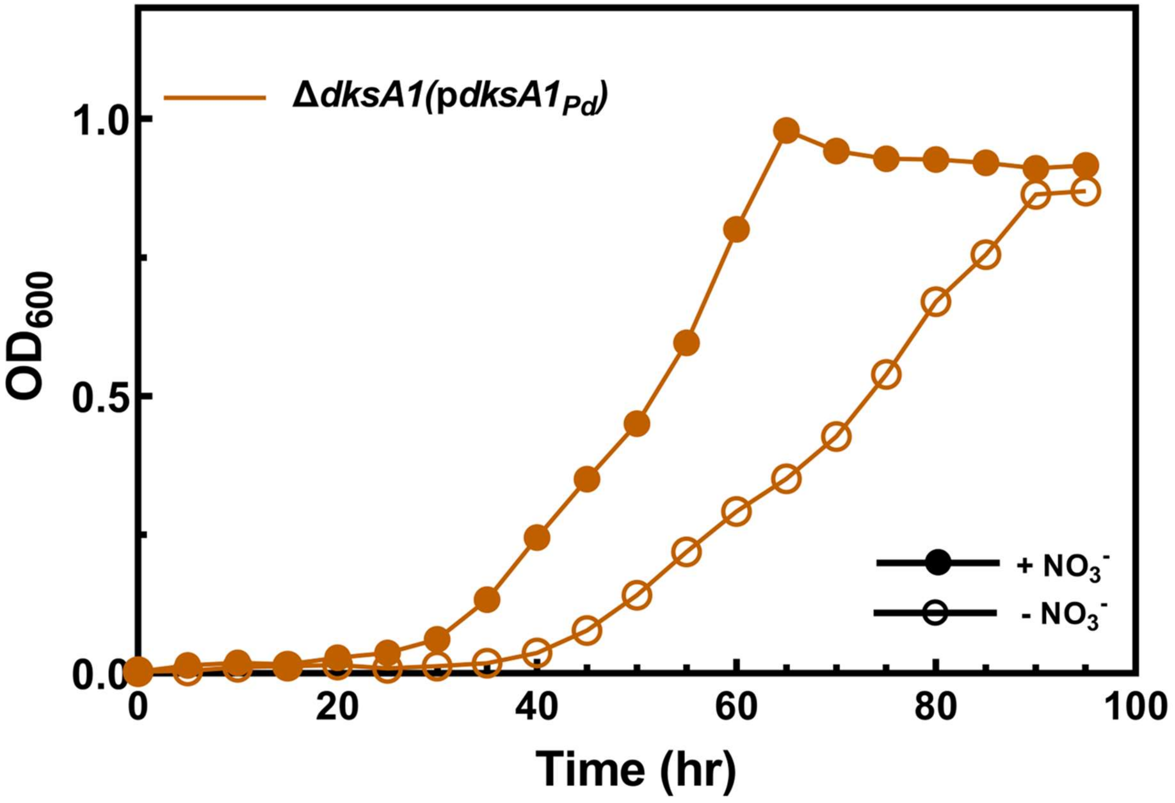


**Figure S4. Complement Δ*dksA1* strain show recovery of the aerobic growth phenotypes.** Aerobic growth of Δ*dksA1*(p*dksA1$_{Pd}$*) in minimal medium with butyrate with (+ $NO_3^-$) or without nitrate (-$NO_3^-$) supplementation. Strains were grown aerobically in minimal medium containing 10 mM butyrate as the sole carbon and energy source with or without 20 mM nitrate supplementation. Absorbance was measured (600 nm) at the indicated time points. The solid line represents the mean of three independent replicates.

### Fig. S5

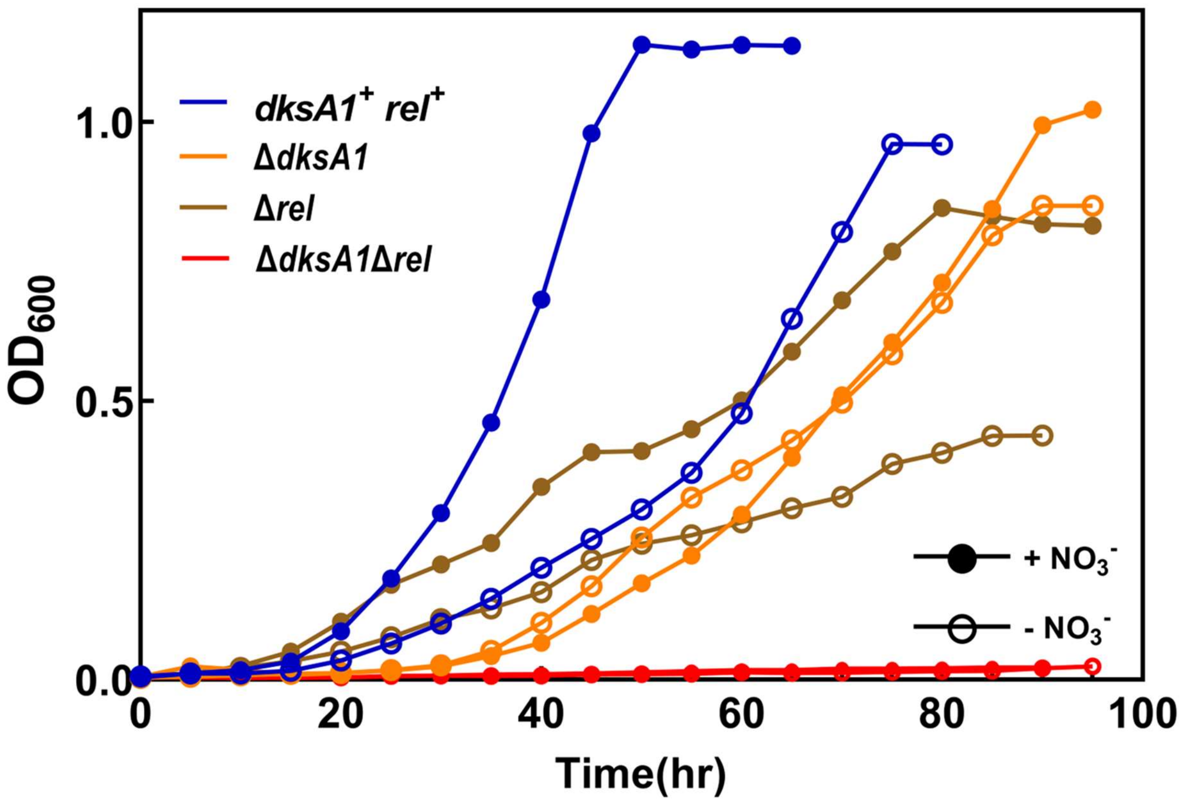


**Figure S5. Combined aerobic growth of indicated strains in butyrate minimal medium with or without nitrate supplementation.** Strains were grown in minimal medium containing 10 mM butyrate as the sole carbon source with (+$NO_3^-$) or without (-$NO_3^-$) 20 mM nitrate supplementation. Absorbance was measured (600 nm) at the indicated time points. The solid line represents the mean of three independent replicates.

### Fig. S6

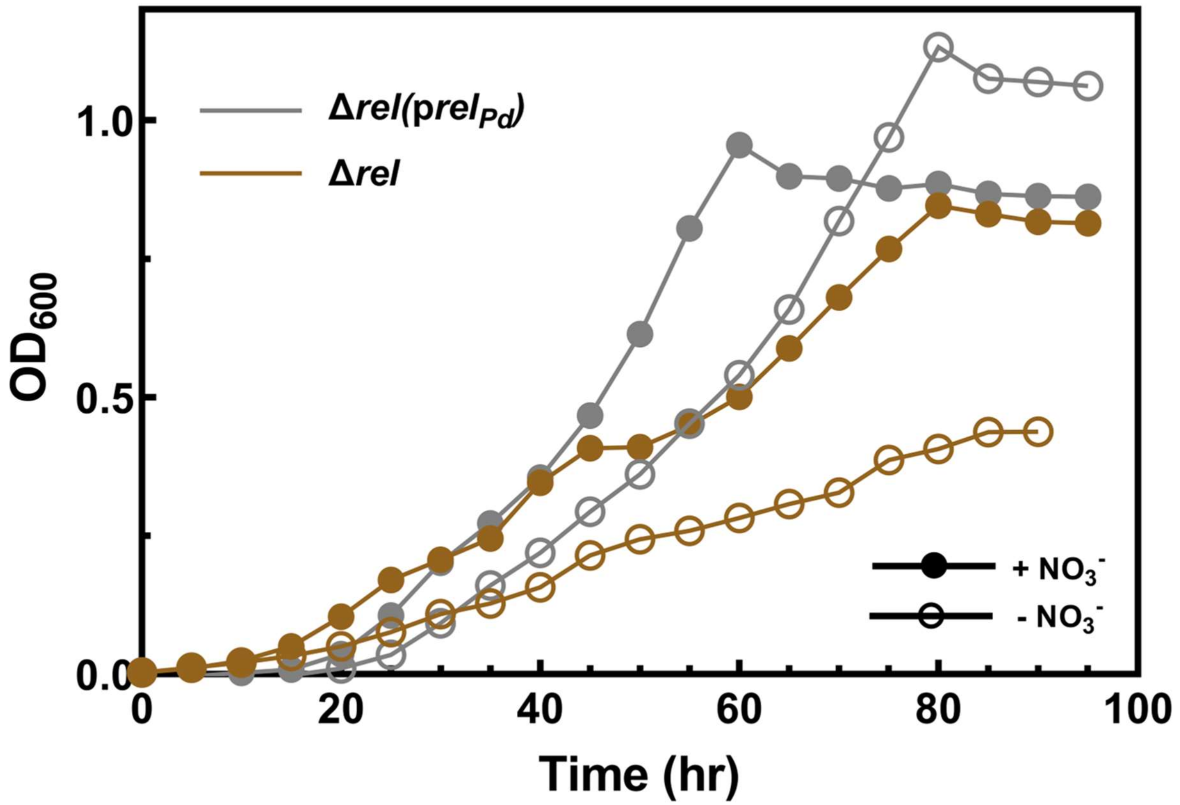


**Figure S6. Complement Δ*rel* strain shows recovery of aerobic growth phenotype.** Aerobic growth of Δ*rel* and Δ*rel*(p*rel$_{Pd}$*) in minimal medium with butyrate with or without nitrate supplementation. Strains were grown aerobically in minimal medium containing 10 mM butyrate as the sole carbon and energy source with or without 20 mM nitrate supplementation. Absorbance was measured (600 nm) at the indicated time points. The solid line represents the mean of three independent replicates.

### Fig. S7

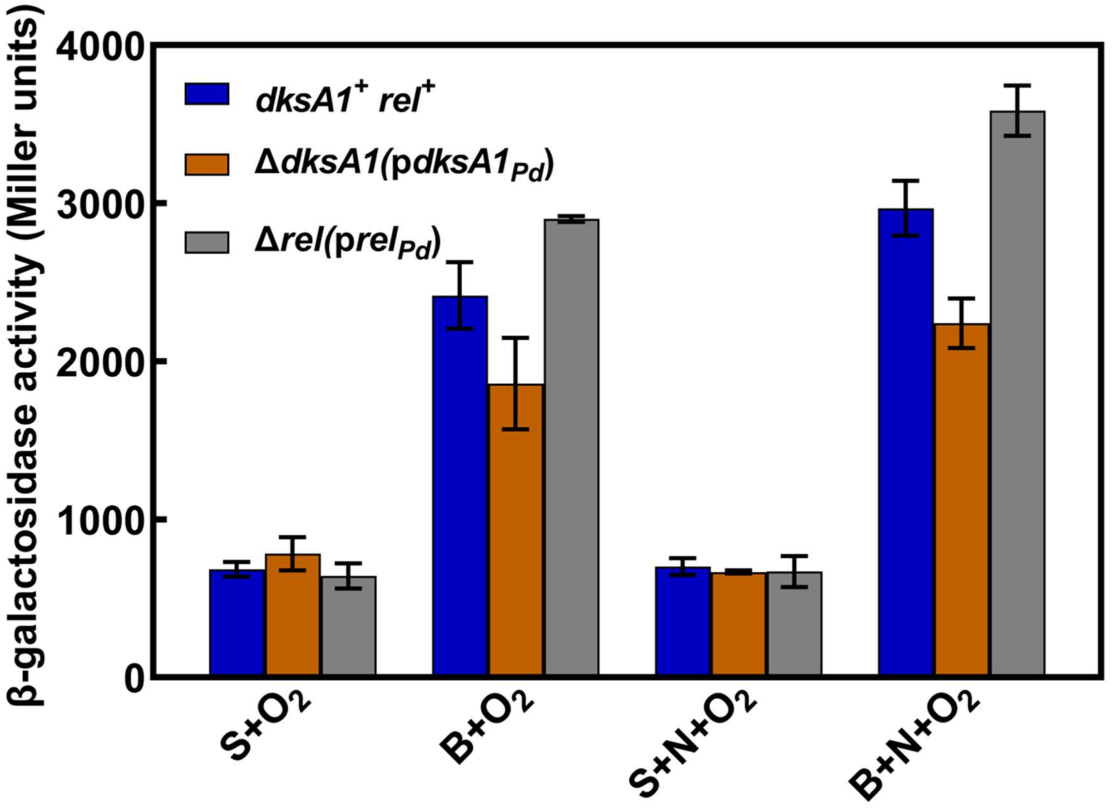


**Figure S7. Complement Δ*dksA1* and Δ*rel* strains show aerobic *nap* promotor activity like wild-type *P. denitrificans*.** β-galactosidase activity in aerobically grown wild-type *P. denitrificans*, Δ*dksA1*(p*dksA1*$_{Pd}$) and Δ*rel*(p*rel*$_{Pd}$) carrying plasmid p*nap-lacZ*. Strains were grown in minimal medium containing 10 mM succinate (S) or butyrate (B) as sole carbon source with or without 20 mM nitrate (N). Aerobic (+$O_2$) cultures were grown in shake flask at 250 rpm. The bars represent the mean of three independent experiments and the error bars represent the standard deviation of the mean.

### Fig. S8

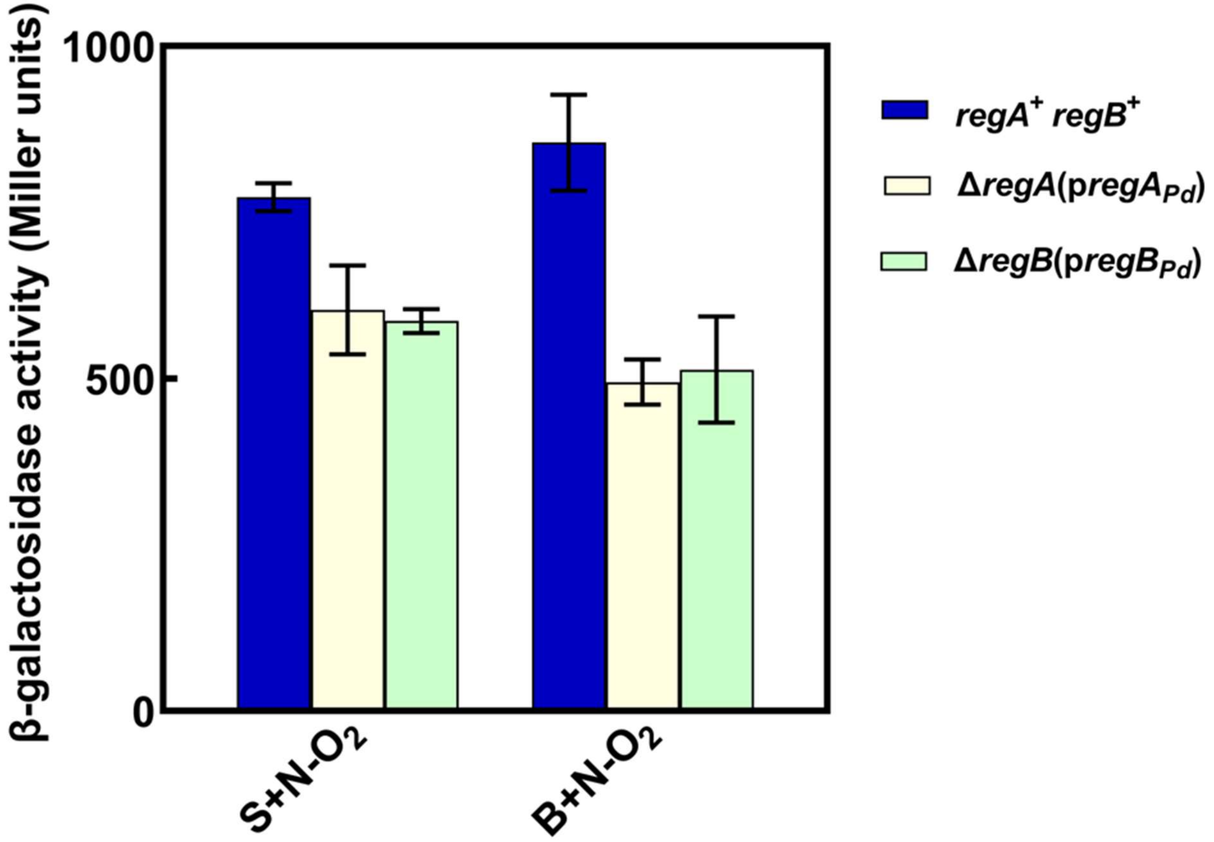


**Figure S8. Complement Δ*regA* and Δ*regB* strains show anaerobic *nap* promotor activity like wild-type *P. denitrificans*.** β-galactosidase activity in anaerobically grown wild-type *P. denitrificans*, Δ*regA*(p*regA*$_{Pd}$) and Δ*regB*(p*regB*$_{Pd}$) carrying plasmid p*nap*-*lacZ*. Strains were grown in minimal medium containing 10 mM succinate (S) or butyrate (B) as sole carbon source with 20 mM nitrate (N). Anaerobic (-$O_2$) cultures were grown in filled closed cap bottles statically. The bars represent the mean of three independent experiments and the error bars represent the standard deviation of the mean.

### Fig. S9

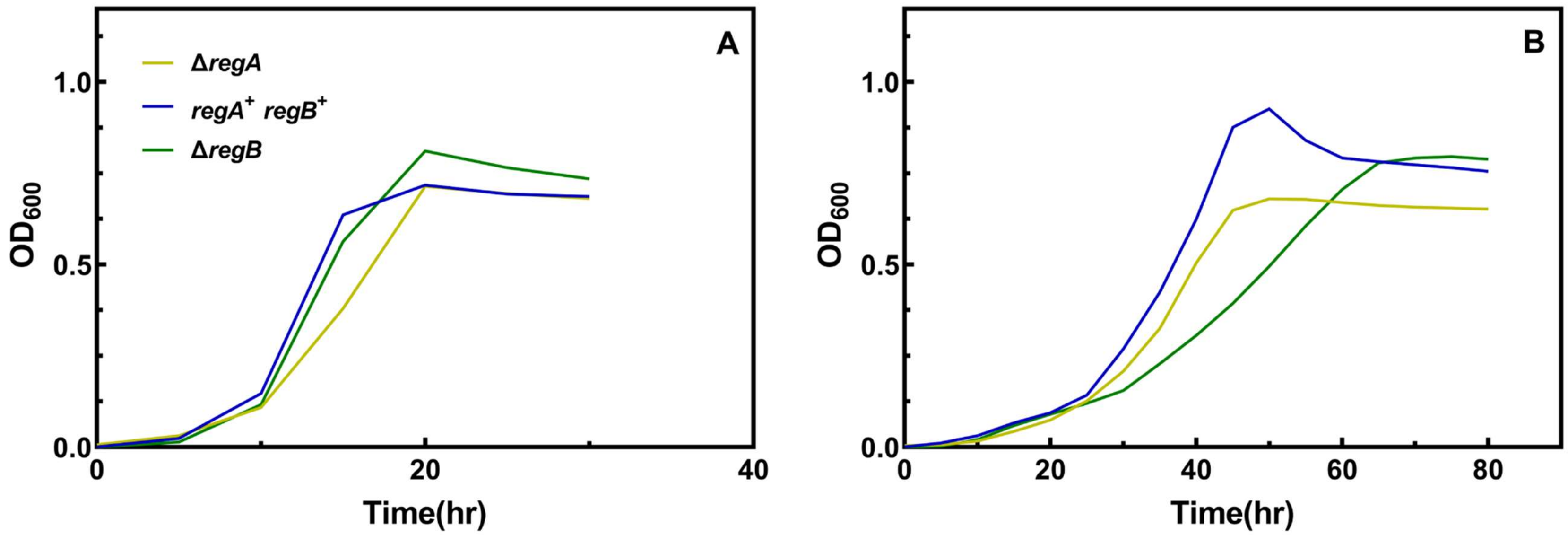


**Figure S9. Δ*regA* strains have reduced anaerobic growth yield.** Anaerobic growth of wild-type *P. denitrificans*, Δ*regA* and Δ*regB* strains in minimal medium containing **(A)** succinate or **(B)** butyrate, with nitrate are shown. Strains were grown anaerobically in minimal medium containing 10 mM succinate or butyrate as the sole carbon and energy source with 20 mM nitrate supplementation. The solid line represents the mean of three independent replicates.

### Fig. S10

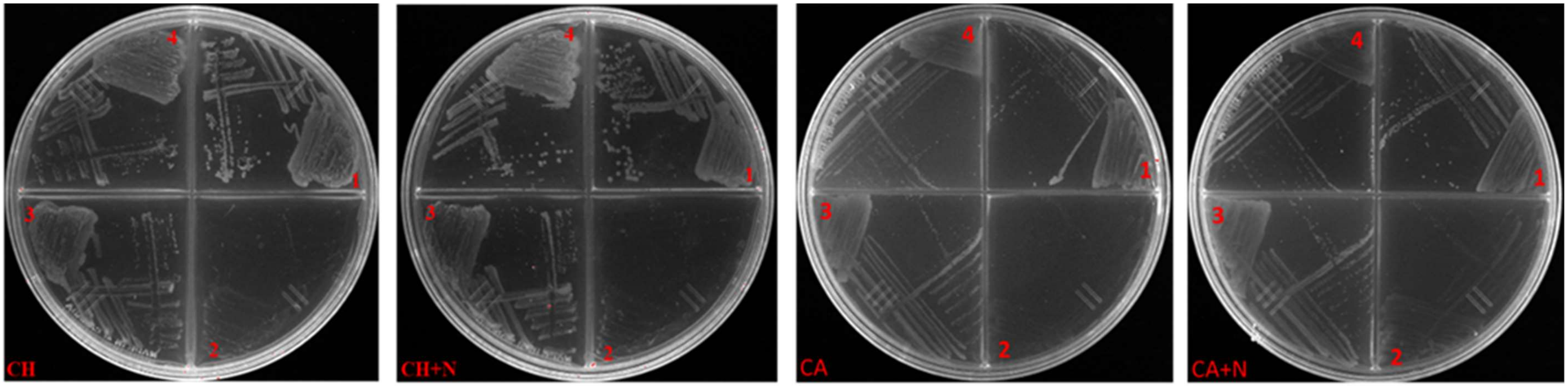


**Figure S10. Δ*dksA1*Δ*rel* is non-viable on reduced carbon substrates**. Aerobic growth of 1 – wild-type *P. denitrificans*; 2 - Δ*dksA1*Δ*rel*; 3 - Δ*dksA1*Δ*rel*(p*dksA1*$_{Pd}$); 4 - Δ*dksA1*Δ*rel*(p*rel*$_{Pd}$) strains on minimal medium plates with choline (CH) or caproate (CA) as the sole carbon source with or without nitrate (N). Strains were streaked on minimal medium agar plates containing 30 mM choline or 7 mM caproate with or without 20 mM nitrate. Plates were incubated overnight at 30 ºC.

### Fig. S11

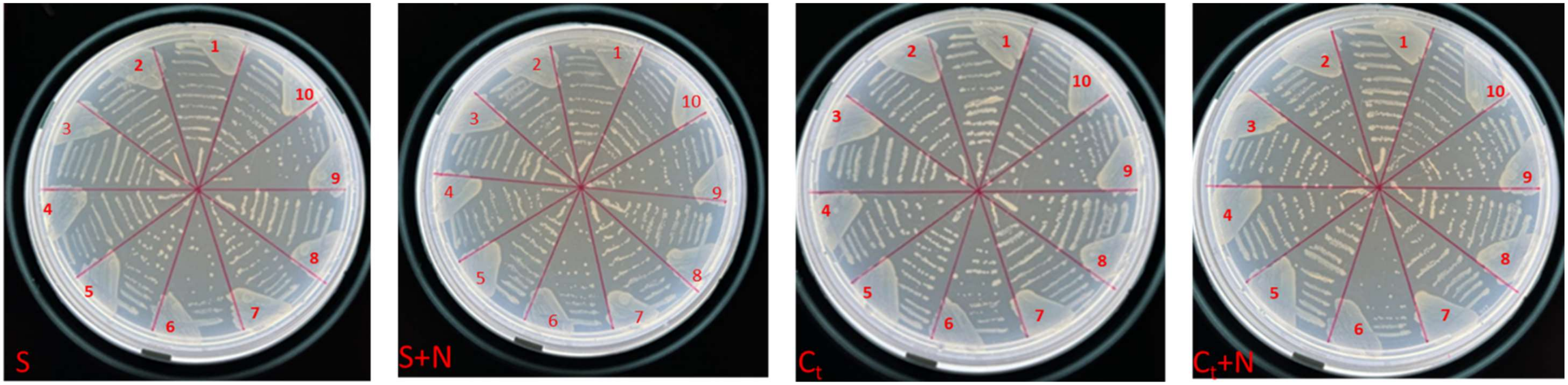


**Figure S11. Δ*dksA1*Δ*rel* is viable on oxidized carbon substrates.** Aerobic growth of 1 - wild-type *P. denitrificans*; 2 – Δ*dksA1*; 3 – Δ*dksA2*; 4 - TM (Δ*dksA1*Δ*dksA2*Δ*Pden_1862*); 5 - Δ*rel* 6 -Δ*dksA1*Δ*rel*; 7 - TMΔ*rel*; 8 - Δ*dksA1*Δ*rel*(p*dksA1*$_{Pd}$); 9 - Δ*dksA1*Δ*rel*(p*rel*$_{Pd}$); 10 - Δ*dksA1*Δ*rel*(p*dksA*$_{Ec}$) strains on minimal medium plates with either succinate (S) or citrate (Ct) as the sole carbon source with or without nitrate (N). Strains were streaked on minimal medium agar plates containing 10 mM succinate or citrate as carbon substrate with or without 20 mM nitrate. Plates were incubated overnight at 30 $^{o}$C.

### Fig. S12

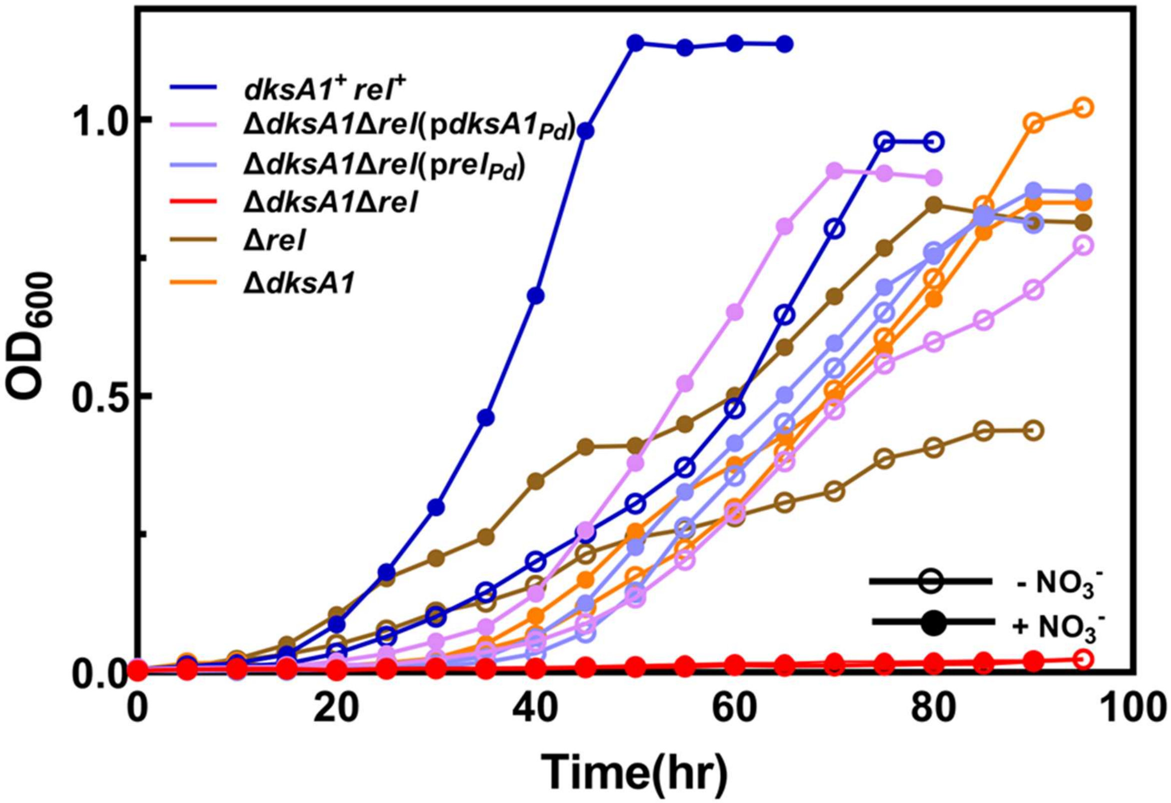


**Figure S12. Combined aerobic growth of mutant and complement strains in butyrate minimal medium with or without nitrate supplementation.** Strains were grown in minimal medium containing 10 mM butyrate as the sole carbon source with (+$NO_3^-$) or without (-$NO_3^-$) 20 mM nitrate supplementation. Absorbance was measured (600 nm) at the indicated time points. The solid line represents the mean of three independent replicates.
