## Supplemental Tables 1-3 for "DksA, ppGpp and RegAB regulate nitrate respiration in *Paracoccus denitrificans*"

### Table S1: Strains and plasmids

| Strain or Plasmid | Relevant genotype or description | Source or Reference |
|---|---|---|
| **<u>Strains</u>** | | |
| ***P. denitrificans*** | | |
| Pd1222 | Rif$^R$, Spec$^R$ | (62) |
| UTD918 | Pd1222 Δ*Pden_0547* (*dksA1*) | This study |
| UTD919 | Pd1222 Δ*Pden_0916* (*dksA2*) | This study |
| UTD920 | Pd1222 Δ*Pden_1862* (*dksA3*) | This study |
| UTD921 | Pd1222 Δ*dksA1*Δ*dksA2* | This study |
| UTD922 | Pd1222 Δ*dksA1*Δ*dksA2*Δ*dksA3* | This study |
| UTD923 | Pd1222 Δ*Pden_1400* (*rel*) | This study |
| UTD924 | Pd1222 Δ*dksA1*Δ*rel* | This study |
| UTD925 | UTD922 Δ*rel* | This study |
| UTD926 | Pd1222 Δ*Pden_2781*::*kan* (*regA*::*kan*) | Maria Torres |
| UTD927 | Pd1222 Δ*Pden_2779* (*regB*) | This study |
| UTD848 | Pd1222 Δ*Pden_1689* (*hmp*) | (6) |
| UTD928 | Pd1222 Δ*dksA2*Δ*hmp* | This study |
| UTD831 | Pd1222 Δ*Pden_1690* (*nsrR*) | (6) |
| *Pd* (p*hmp*) | *Pd*1222 + p*hmp-lacZ* | (6) |
| UTD918(p*hmp*) | UTD918 + p*hmp-lacZ* | This study |
| UTD919(p*hmp*) | UTD919 + p*hmp-lacZ* | This study |
| UTD922(p*hmp*) | UTD922 + p*hmp-lacZ* | This study |
| UTD923(p*hmp*) | UTD923 + p*hmp-lacZ* | This study |
| *Pd* (p*nap*) | *Pd*1222 + p*nap-lacZ* | This study |
| UTD918(p*nap*) | UTD918 + p*nap-lacZ* | This study |
| UTD919(p*nap*) | UTD919 + p*nap-lacZ* | This study |
| UTD922(p*nap*) | UTD922 + p*nap-lacZ* | This study |
| UTD923(p*nap*) | UTD923 + p*nap-lacZ* | This study |
| Δ*dksA1*(p*dksA1*$_{Pd}$) | Δ*dksA1* + pIND4-*dksA1* | This study |
| Δ*dksA2*(p*dksA2*$_{Pd}$) | Δ*dksA2* + pIND4-*dksA2* | This study |
| Δ*rel*(p*rel*$_{Pd}$) | Δ*rel* + pIND4-*rel* | This study |
| Δ*regA*(p*regA*$_{Pd}$) | Δ*regA* + pIND4-*regA* | This study |
| Δ*regB*(p*regB*$_{Pd}$) | Δ*regB* + pIND4-*regB* | This study |
| Δ*dksA1*Δ*rel*(p*dksA1*$_{Pd}$) | Δ*dksA1*Δ*rel* + pIND4-*dksA1* | This study |
| Δ*dksA1*Δ*rel*(p*rel*$_{Pd}$) | Δ*dksA1*Δ*rel* + pIND4-*rel* | This study |
| ***E. coli*** | | |
| DH5α | *supE44*, Δ*lacU169* (φ80 *lacZ*ΔM15), *hsdR1*, *recA1*, *endA1*, *gyrA96*, *thi-1*, *relA1*, pRK2013 | (64) |
| S17-1 | *pro*, $res^-$, $mod^+$, *recA* derivative of EC294 with integrated RP4-2 [*Tc::Mu*][*Km::Tn7*], Tp$^R$ | (63) |
| MG1655 | Wild-type, *rph-1* | Genetic Stock Center |
| Δ*dksA*$_{Ec}$ | *E. coli* K-12 BW25113 Δ*dksA* | KEIO library |
| Δ*dksA*$_{Ec}$(p*dksA1*$_{Pd}$) | Δ*dksA*$_{Ec}$ + pIND4-*dksA1* | This study |
| Δ*dksA*$_{Ec}$(p*dksA2*$_{Pd}$) | Δ*dksA*$_{Ec}$ + pIND4-*dksA2* | This study |
| **<u>Plasmids</u>** | | |
| pK18*mobsacB* | pMB1, *mob*, *nptII*, *sacB*, Suc$^S$, Km$^R$ | (67) |
| pST-Blue1 | SP6/T7 promoters, Amp$^R$, Km$^R$ | Novagen |
| pRK2013 | ColE1 *ori*, $mob^+$, $tra^+$, Km$^R$ | (64) |
| pMP220 | IncP replicon, Broad-host-range, low-copy-number *lacZ* promoter-fusion vector, Tc$^R$ | (70) |
| p*hmp-lacZ* | pMP220 *hmpp-lacZ* transcriptional fusion, Tc$^R$ | (6) |
| p*nap-lacZ* | pMP220 *napp-lacZ* transcriptional fusion, Tc$^R$ | This study |
| pIND4 | ColE1 *ori*, pMG160 *ori*, $lacI^q$, Km$^R$ | (71) |
| pIND4-*dksA1* | pIND4 Ω[BamHI: *P.den dksA1*] | This study |
| pIND4-*dksA2* | pIND4 Ω[BamHI: *P.den dksA2*] | This study |
| pIND4-*rel* | pIND4 Ω[BamHI: *P.den rel*] | This study |
| pIND4-*regA* | pIND4 Ω[BamHI: *P.den regA*] | This study |
| pIND4-*regB* | pIND4 Ω[BamHI: *P.den regB*] | This study |

#### Table S2: List of primers

| Primer name | Primer Sequence (5' to 3') |
|---|---|
| **For deletion** | |
| *dksA1*-R1 | *GGAATTCATGAAGACCTGCACCATGGGCG* |
| *dksA1*-R1 | *ACGATGAAAGTGCATCGCGACGACTGAACG* |
| *dksA1*-F2 | *GCGATGCACTTTCATCGTTCCTCGGGGAGCC* |
| *dksA1*-R2 | *AACTGCAGCAGCTTGCCCTTGCGGATGTAG* |
| *dksA1-F3* | *CATGAACGAGCGGCAACTCG* |
| *dksA1*-R3 | *TGCGCTTCCAGCGTCATGGT* |
| *dksA2*-F1 | *GGAATTCGCAATCACCGTGCAGCCGGT* |
| *dksA2-R*1 | *GTGTCAGAACCTACATGCCTGCCTGGGCTT* |
| *dksA2*-F2 | *GCATGTAGGTTCTGACACGCCTGGACGGTG* |
| *dksA2-R2* | *AACTGCAGCTGCTCAATAAGGTGCCGGCG* |
| *dksA2*-F3 | *GAAACCCTTGACGTCTGGGAA* |
| *dksA2-R3* | *TGGCGGCAAAGCCTAAACCAG* |
| *dksA3*-F1 | *AACTGCAGCCACCCGATCCTTCATCAG* |
| *dksA3*-R1 | *GCTCTAGAGTCAGCTCCGTTCCGCCTGCC* |
| *dksA3*-F2 | *GCTCTAGAGGGCTTGCCTACCAGCGCAG* |
| *dksA3*-R2 | *CGGAATTCGAACAGCGACGGGCAATATCG* |
| *dksA3*-F3 | *CTTCTGCGAGGAATGCGACGAG* |
| *rel*-F1 | *CGGGATCCTCGCCGTCAGCAGATTC* |
| *rel*-R1 | *GCTCTAGACTTCCAGCATCATGCGC* |
| *rel*-F2 | *GCTCTAGAAGCTATGCTCAGATGTGGAC* |
| *rel*-R2 | *CCCAAGCTTCGATCACCATGAACCGGA* |
| *rel*-F3 | *ACGGCTTTCGCATCATCAC* |
| *rel*-R3 | *GATAGGTGATGCCGACGATG* |
| *regB*-F1 | *AACTGCAGAGCCGGGAATCACAAGATAGG* |
| *regB*-R1 | *GCTCTAGAGCTGGGTCTTTCTGTTGGG* |
| *regB*-F2 | *GCTCTAGAATCCGACTGTATCGTTCAGG* |
| *regB*-R2 | *CCCAAGCTTCTCGGCCAGGTTCAGATAG* |
| *regB*-F3 | *CAGCCTTCTGCTGGTCGATG* |
| *regB*-R3 | *TAACGCGGGCTGCGTTTG* |
| **For promotor fusion** | |
| *napp*-F | *CGGAATTCCCGGCTTGACTTGCCATAC* |
| *napp*-R | *AACTGCAGCATGGCCGCTCTCTAATGC* |
| **For complementation** | |
| pIND4-F | *TCACCAGCTCACCGTCTTTC* |
| pIND4-R | *ACGTAA ATG CAT GCC GCT TC* |
| pIND4-*dksA1*-F | *GGTGATGGTGATGAGATCTGATGAAAGCCCAGACCTTTCTG* |
| pIND4-*dksA1-R* | *GGAGAAATTAACCATGGGAGTCAGTCGTCGCGATGCAC* |
| pIND4-*dksA2*-F | *AGCTAATTAAGCTTAGTGATGGTGATGGTGATGAGATCTGGGCATGTAGGCCGGAAAATC* |
| pIND4-*dksA2*-R | *CACACATCTAGAATTAAAGAGGAGAAATTAACCATGGGAGATCCATGGATCCGGTTGC* |
| pIND4-*dksA*$_{Ec}$-F | *AGCTAATTAAGCTTAGTGATGGTGATGGTGATGAGATCTGGTAAACGTGATGGAACGG* |
| pIND4-*dksA*$_{Ec}$-R | *CACACATCTAGAATTAAAGAGGAGAAATTAACCATGGGAGATGCAAGAAGGGCAAAAC* |
| pIND4-*rel*-F | *AGCTAATTAAGCTTAGTGATGGTGATGGTGATGAGATCTGTCAGTTCATATGGGCGGAAGG* |
| pIND4-*rel*-R | *CACACATCTAGAATTAAAGAGGAGAAATTAACCATGGGAGGGACGATCTGCGCGAGCG* |
| pIND4-*regA*-F | *GCTTAGTGATGGTGATGGTGATGAGATCTGATGAGCGAATGATCGGATTGACC* |
| pIND4-*regA*-R | *GAATTAAAGAGGAGAAATTAACCATGGGAGTTAACGCGGGCTGCGTTTGG* |
| pIND4-*regB*-F | *GTGATGGTGATGGTGATGAGATCTGGACATCCGGGCAATAGGTATATCC* |
| pIND4-*regB*-R | *AAAGAGGAGAAATTAACCATGGGAGTCAGGCCAGTATCTCGGGGT* |

#### Table S3A: Doubling time of strains during aerobic growth in minimal medium

| | | Doubling Time (hours) | | | | | |
|---|---|---|---|---|---|---|---|
| **Medium Carbon Source** | **Growth Condition \ Strain** | ***dksA1⁺rel⁺*** | **Δ*dksA1*** | **Δ*rel*** | **Δ*dksA1*Δ*rel*** | **Δ*dksA1*Δ*rel* (p*dksA1*$_{Pd}$)** | **Δ*dksA1*Δ*rel* (p*rel*$_{Pd}$)** |
| Succinate | Aerobic no $NO_3^-$ | 3.0 ± 0.6 | 3.0 ± 0.5 | n.d* | n.d* | n.d* | n.d* |
| | Aerobic + $NO_3^-$ | 2.6 ± 0.3 | 2.8 ± 0.3 | n.d* | n.d* | n.d* | n.d* |
| Butyrate | Aerobic no $NO_3^-$ | 15.7 ± 0.4 | 18.8 ± 4.1 | 26.1 ± 3.9 | - | 16.5 ± 3.8 | 18.3 ± 3.1 |
| | Aerobic + $NO_3^-$ | 8.0 ± 0.8 | 19.0 ± 2.5 | 17.0± 4.9 | - | 14.2 ± 2.4 | 17.5 ± 2.4 |

*n.d – not determined

#### Table S3B: Doubling time of strains during aerobic growth in minimal medium with or without SHX

| | | Doubling times (hours) | | |
|---|---|---|---|---|
| **Medium Carbon Source** | **Growth Condition \ Strain** | ***dksA1⁺rel⁺*** | **Δ*rel*** | **Δ*rel*(p*rel*$_{Pd}$)** |
| Succinate | Aerobic No SHX | 3.9 ± 0.1 | 4.7 ± 1.3 | 3.4 ± 1.2 |
| | Aerobic 3 mM SHX | 4.7 ± 1.0 | - | 4.2 ± 1.6 |
| | Aerobic 3 mM + 0.2% CAA | 2.8 ± 0.7 | 3.8 ± 1.3 | 2.6 ± 1.3 |

SHX – Serine hydroxamate; CAA – Casamino acids
